# Sex-dimorphic effects of neuromelanin buildup in rodent nigral dopamine neurons: implications for sex-biased vulnerability in Parkinson’s disease

**DOI:** 10.1101/2025.04.18.649338

**Authors:** Sebastian Luca D’Addario, Mariangela Massaro Cenere, Francesca Cossa, Silvia Scaricamazza, Valentina Nesci, Ginevra D’Ottavio, Daniele Caprioli, Alberto Ferri, Miquel Vila, Ada Ledonne, Nicola Biagio Mercuri

## Abstract

Neuromelanin (NM) is a dark pigment accumulating with age in human substantia nigra pars compacta (SNpc) dopamine (DA) neurons, conferring the dark look that inspired nigral area’s name. Despite NM has long been associated with Parkinson’s disease (PD), as melanized neurons favorably degenerate during disease development, NM functions within SNpc DA neurons are still mostly elusive. Here, by exploiting an NM-producing rat model generated by viral vector-induced expression of human Tyrosinase (hTyr), we inspected NM impact on nigral DA neurons’ survival and activity, on mitochondrial functionality of SNpc, and behaviors resembling non-motor and motor PD symptoms. Our data reveal sex dimorphism in NM effects on nigrostriatal dopamine circuit, with sex-biased alterations in neuronal firing activity and underlying intrinsic currents, nigral mitochondrial functions, and non-motor PD symptoms (anxiety). In conclusion, this study discloses unrealized NM effects within nigral DA neurons, advancing our comprehension of sex-specific features shaping sex-biased vulnerability to PD.

## Introduction

Parkinson’s disease (PD) is characterized by a multifaceted symptomatology including motor anomalies (rigidity, resting tremor, postural instability, and bradykinesia) and a spectrum of non-motor symptoms, counting sleep disturbances, autonomic dysfunctions, cognitive deficits, psychiatric manifestations like depression, anxiety, and apathy associated to prodromal stages, and psychosis and dementia arising at late diseases stages^1,2,3^. Although PD diagnosis relies on appearance of cardinal motor deficits, it is now recognized that prodromal non-motor symptoms can precede them of several years^2^. Thus, disclosure of mechanisms instrumental to both motor and non-motor symptoms is key to comprehending PD etiology and progression.

Increasing evidence indicates that biological sex is an important factor influencing PD development and phenotypic expression^4^. The risk of developing PD is higher in men than in women, with an estimated global prevalence of 1.4:1 for males versus females. Nonetheless, women manifest a faster pathological progression, associated with a higher mortality rate. Sex-related differences also involve symptomatology - with some symptoms being more prevalent in one sex than the other - and response to pharmacological and surgical PD therapy^4,5^. Despite this extensive evidence, the investigation of sex dimorphism in PD animal models remains mostly neglected, failing to provide essential information to understand processes governing sex-related vulnerability in PD patients^4,5^.

The degeneration of dopamine (DA) neurons of the substantia nigra pars compacta (SNpc) is the main hallmark of PD. While it is recognized that multiple pathological processes can undermine nigral DA neuron viability (i.e., misfolded α-synuclein (α-syn) aggregation, oxidative stress, mitochondrial dysfunction, impairment of the ubiquitin-proteasome system, and neuroinflammation), the precise cascade of events liable for their degeneration in PD remains unidentified^2^. A key part could be played by neuromelanin (NM)^6^, a dark brown cytoplasmatic pigment naturally produced in human catecholaminergic neurons with highest accumulation in SNpc DA neurons. NM appears around the third year of life in human SNpc and progressively accumulates with age^7^ conferring the dark appearance that inspired nigral area’s nomenclature. NM is a complex polymeric molecule, made of a melanin structure bound with lipids, peptides, and metals^8^; it is synthesized as by-product of DA metabolism, with non-enzymatic DA autoxidation led by ferric ion producing quinones, that by multiple steps of polymerization and aggregation with proteins, lipids, and metals constitute the final NM pigment^9^. Beyond DA autoxidation, NM biosynthesis may rely on an enzymatic pathway lead by Tyrosinase (Tyr) starting from tyrosine oxidation. Tyr is responsible for the production of peripheral melanins in skin and hairs^10^, while due to limited protein amounts detected in human SNpc^11,12,13^, its contribution to NM production remains still debated.

A causal relationship between NM buildup and PD pathogenesis has been inferred based on earlier evidence that NM-containing neurons preferentially degenerate during PD progression^14^ and intracellular inclusions resembling Lewy bodies (LBs) mainly form in proximity to NM granules^15^. Later advances on understanding of NM functions in nigral DA neurons have been limited by the absence of adequate preclinical models, as NM is not endogenously produced in rodents, opposite to humans. The recent introduction of first NM-producing rodent models, generated by inducing overexpression of human Tyr (hTyr) by adeno-associated viral vector (AAV) or genetic manipulations, provided direct evidence on NM-dependent histopathological alterations in the nigrostriatal DA circuits, corroborating the idea that abnormal NM levels contribute to PD development^16,17,18,19^. Nevertheless, multiple aspects inherent to NM accumulation within nigral DA neurons are still enigmatic: it is unknown whether (and how) NM affects SNpc DA neurons’ activity, if the vulnerability of nigral DA neurons to NM buildup is sex-biased, and there is sex dimorphism in NM roles in promoting nonmotor and motor PD symptoms. To increase knowledge of these aspects, in this study we generated a novel NM-producing rat model by inducing AAV-dependent hTyr expression in the SNpc. We provide original evidence on sex-biased NM roles in shaping nigral DA neurons’ functions and fostering emergence of non-motor PD symptoms, thus gaining insight into sex-specific processes that could influence sex-biased vulnerability to PD development.

## Results

### AAV-induced hTyr expression triggers NM production in nigral DA neurons

To evaluate NM-induced effects on nigral DA neurons, we generated a model of hTyr overexpression based on bilateral injection of serotype 9 AAV carrying hTyr (AAV-hTyr) above the SNpc of rats of both sexes (**Fig. 1A**). After 4 weeks, NM production - visible to the naked eye as a dark brown area entirely covering the SNpc - appeared in AAV-hTyr-injected rats but not in control animals inoculated with an AAV empty vector (AAV-null) (**Fig. 1B**). Quantification of hTyr by immunofluorescence (**Fig. 1C**) revealed a similar number of hTyr-expressing nigral DA neurons among sexes (**Fig. 1D**). hTyr-positive DA neurons were also observed in the ventral tegmental area (VTA) (**Extended Data Fig. 1A**). Masson-Fontana staining confirmed NM accumulation at cellular level in SNpc (**Fig. 1E**) and VTA neurons **(Extended Data Fig. 1B)**. Patchy pigmentation also appeared in substantia nigra pars reticulata (SNpr), despite confined to cells with typical morphology of DA neurons (**Extended Data Fig. 1B**). To assess whether there is sex-biased proneness in NM accumulation, we measured NM levels within SNpc DA neurons in female- and male AAV-hTyr-injected rats, reporting higher intracellular NM levels in males. (**Fig. 1F**). Collectively, these findings demonstrate fast NM production by AAV9-induced hTyr overexpression in rats, with a slight sex dimorphism in intracellular NM buildup.

**Figure 1.**
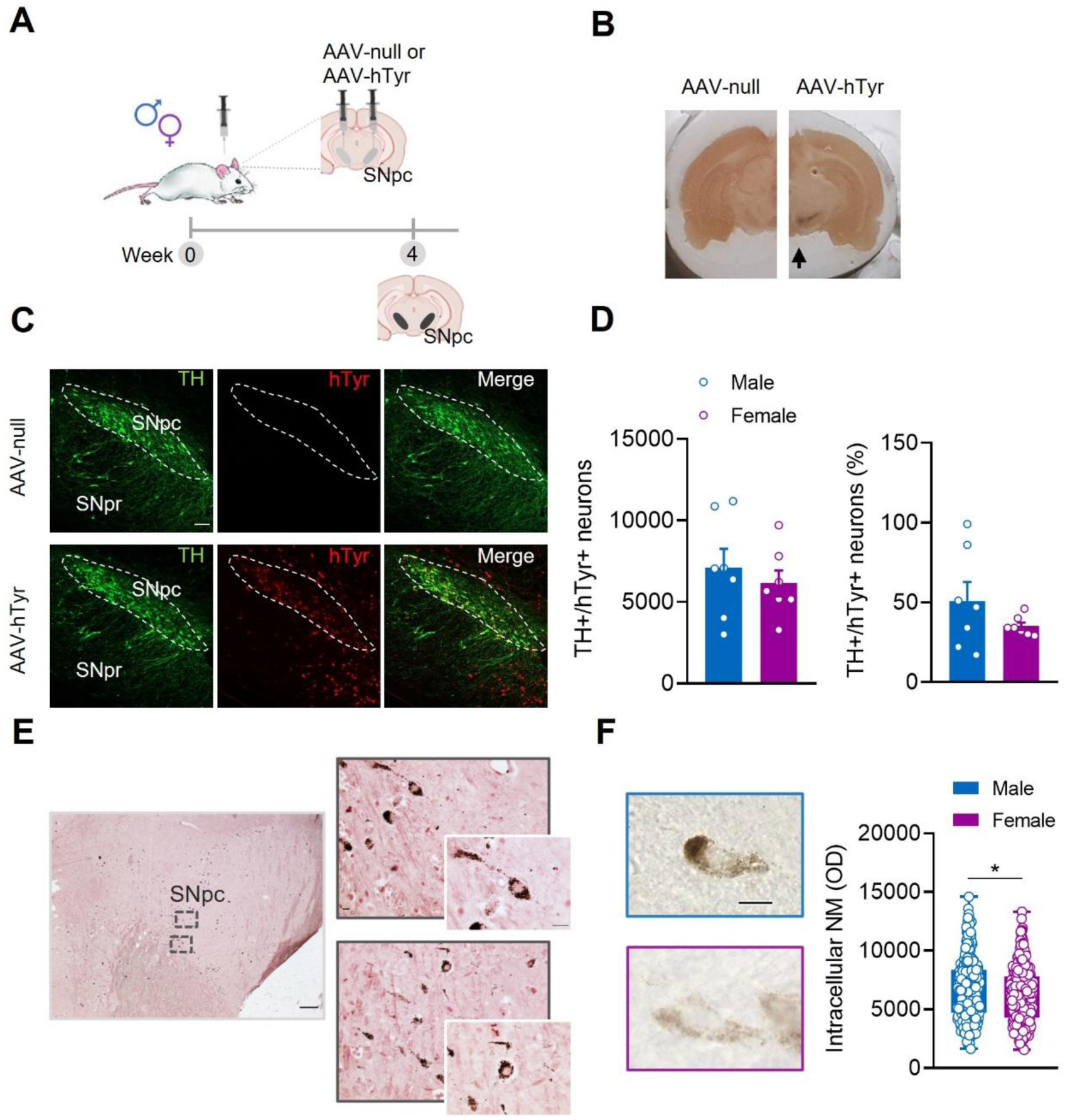
NM production within nigral DA neurons by local AAV-induced human tyrosinase (hTyr) expression **A,** Scheme of the experimental design and timeline used to induce *in vivo* NM production within nigral DA neurons of female and male rats. **B,** Representative unstained rat brains mounted on a cryostat, showing that a dark brown area entirely covering the SNpc (black arrow) can be macroscopically detected in AAV-hTyr-injected rats, but not in control AAV-null animals, four weeks post-AAVs injection. **C,** Representative double-labelled confocal images of coronal SNpc sections immunostained for TH (green) and hTyr (red) (scale bar: 100 μm). **D,** Quantification of hTyr-expressing neurons in male and female AAV-hTyr-injected rats, measured as total number of TH+hTyr+ neurons (left; male *n*=7; female *n*= 7; Student’s *t*-test, P = 0.517) and as percentage of hTyr+ neurons over total TH+ neurons (right; male *n* = 7; female *n* = 7; Student’s *t*-test, P = 0.211). **E,** Representative micrographs of Masson-Fontana staining (showing NM as dark brown granules) in 5-μm-thick SNpc sections from an AAV-hTyr-injected rat (scale bar: 200 μm, 20 μm, and 10 μm). **F,** Representative images of NM-containing neurons from unstained SNpc sections (left; scale bar: 10 μm), and quantification of intracellular NM optical density in male and female AAV-hTyr-injected rats (right; Mann-Whitney test; P = 0.044). Data are expressed as mean ± SEM. **(D, F)**; dots represent individual animals **(D)** or individual cells **(F)**. *P < 0.05.

### NM effects on the nigrostriatal dopamine circuit

To disclose whether nigral NM buildup induces early histopathological alterations in the nigrostriatal DA circuit, first we performed stereological cell counts of SNpc tyrosine hydroxylase positive (TH+) neurons (**Fig. 2A**). We found a reduction in SNpc TH+ cells in male AAV-hTyr-injected rats compared to controls (**Fig. 2B**). To decipher if this relies on TH down-regulation (a typical adaptation of nigral DA neurons to toxic insults) or real neuronal degeneration, we quantified total DA neurons, that is TH+ plus TH negative/NM positive (TH-NM+) cells. We found a slight, but not significant, decrease of total DA neurons, implying that NM at this stage does not cause substantial neuronal death, but just TH down-regulation (**Fig. 2C**). Likewise, females showed fewer TH+ cells (**Fig. 2D**), but unaltered total DA neurons (**Fig. 2E**). Next, we evaluated NM-induced effects on VTA DA neurons, observing equal number of TH+ neurons and total DA neurons either in male-(**Fig. 2F-G**) and female (**Fig. 2I-J**) AAV-hTyr-injected rats and respective controls. The percentage of NM-producing cells was analogous between SNpc and VTA in both sexes (**Fig. 2H-K**), further supporting a preferential vulnerability of SNpc (over VTA) DA neurons to NM buildup.

**Figure 2.**
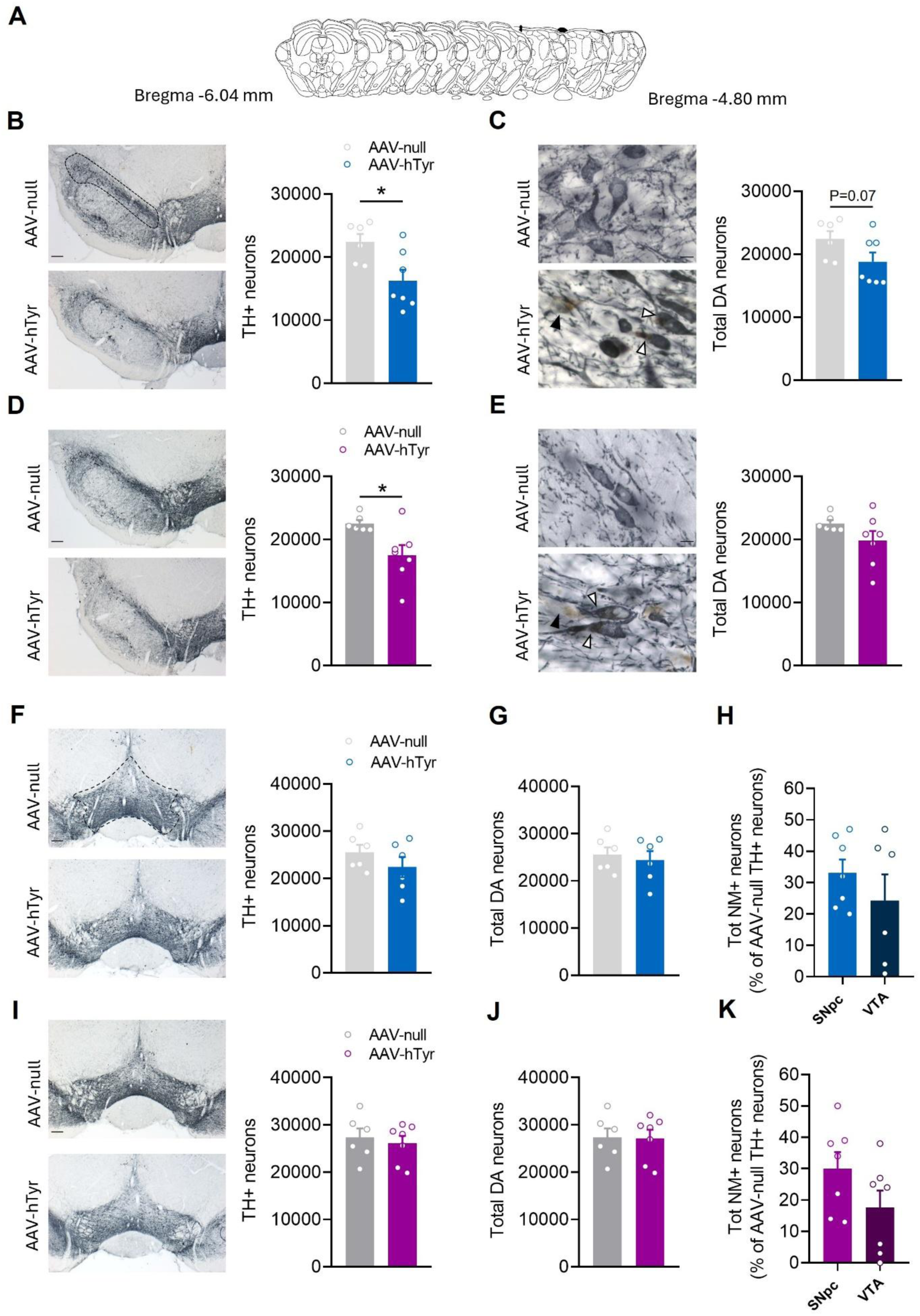
Nigral NM buildup downregulates tyrosine hydroxylase in SNpc DA neurons independently from sex. **A,** Qualitative mouse brain atlas showing selected SNpc serial sections for quantitative histological analysis. **B,** Representative SNpc images (left, scale bar: 200 μm), and quantification (right) of TH+ neurons in male rats (AAV-null *n* = 6; AAV-hTyr *n* = 7; Student’s *t*-test, P = 0.018). **C,** TH-immunostained SNpc showing TH−NM+ (black arrowhead) and TH+NM+ (white arrowhead) neurons (left, scale bar: 10μm), and histogram (right) of total DA neurons in SNpc of male rats (AAV-null *n* = 6; AAV-hTyr *n* = 7; Mann-Whitney test, P = 0.073). **D,** Representative SNpc images (left, scale bar: 200 μm), and quantification (right) of TH+ neurons in female rats (AAV-null *n* = 6; AAV-hTyr *n* = 7; Student’s *t*-test with Welch’s correction, P = 0.020). **E,** TH-immunostained SNpc showing TH−NM+ (black arrowhead) and TH+NM+ (white arrowhead) neurons (left, scale bar: 10 μm), and histogram (right) of total SNpc DA neurons of female rats (AAV-null *n* = 6; AAV-hTyr *n* = 7; Student’s *t*-test with Welch’s correction, P = 0.135). **F,** Representative VTA images (left, scale bar: 200 μm), and quantification (right) of TH+ neurons in male rats (AAV-null *n* = 6; AAV-hTyr *n* = 6; Student’s *t*-test, P = 0.261). **G,** Count of total VTA DA neurons of male rats (AAV-null *n* = 6; AAV-hTyr *n* = 6; Student’s *t*-test, P = 0.656). **H,** Quantification of total NM-containing neurons (as % of AAV-null TH-immunopositive neurons) in the SNpc and VTA of male AAV-hTyr rats (SNpc *n* = 6; VTA *n* = 6; Student’s *t*-test, P = 0.344). **I,** Representative VTA images (left, scale bar: 200 μm), and quantification (right) of TH+ neurons in female rats (AAV-null *n* = 6; AAV-hTyr *n* = 7; Student’s *t*-test, P = 0.640). **J,** Count of total VTA DA neurons of female rats (AAV-null *n* = 6; AAV-hTyr *n* = 7; Student’s *t*-test, P = 0.948). **K,** Quantification of total NM-containing neurons (as % of AAV-null TH-immunopositive neurons) in the SNpc and VTA of female AAV-hTyr rats (SNpc *n* = 6; VTA *n* = 6; Student’s *t*-test, P = 0.128). Data are expressed as mean ± SEM **(B-K)**; dots represent individual animals. *P < 0.05.

Moreover, we quantified TH density in the dorsal striatum (dST) in three fields along the rostrocaudal axes (**Extended Data Fig. 2A-C**) reporting regular levels both in males (**Extended Data Fig. 2D-F**) and females (**Extended Data Fig. 2G-I**). Analyses of TH density in the ventral striatum (i.e., nucleus accumbens (NAc) core and NAc shell (dorsal and ventral parts) proved male AAV-hTyr-injected rats have regular TH levels in NAc core **(Extended Data Fig. 3A)** and ventral NAc shell **(Extended Data Fig. 3C),** and a slight, but not statistically significant, reduction in the dorsal NAc shell **(Extended Data Fig. 3B)**. Female AAV-hTyr-injected rats displayed unaltered TH density in the analyzed NAc parts **(Extended Data Fig. 3D-F).**

### NM accumulation affects the mitochondrial function in SNpc in a sex-specific manner

Mitochondria are key to proper cellular bioenergetics and their dysfunction contributes to PD development^20^. To assess whether NM overload damages mitochondrial functionality, we evaluated mitochondrial respiration in freshly dissected SNpc punches from AAV-hTyr-injected rats and controls (**Fig. 3A**). Real-time measurement of oxygen consumption rate (OCR) in male NM-producing rats (**Fig. 3B**) revealed unaltered basal respiration and ATP production but increased maximal respiration and spare capacity (**Fig. 3C**). Thus, NM buildup in males appears to strain mitochondrial function. Differently, NM overload did not affect nigral mitochondrial functionality in female AAV-hTyr-injected rats, whose OCR (**Fig. 3D**), basal and maximal respiration, ATP production, and spare capacity (**Fig. 3E**) are comparable to the control. These data prove that NM load within SNpc DA neurons promotes sex-dimorphic nigral mitochondrial adaptations.

**Figure 3.**
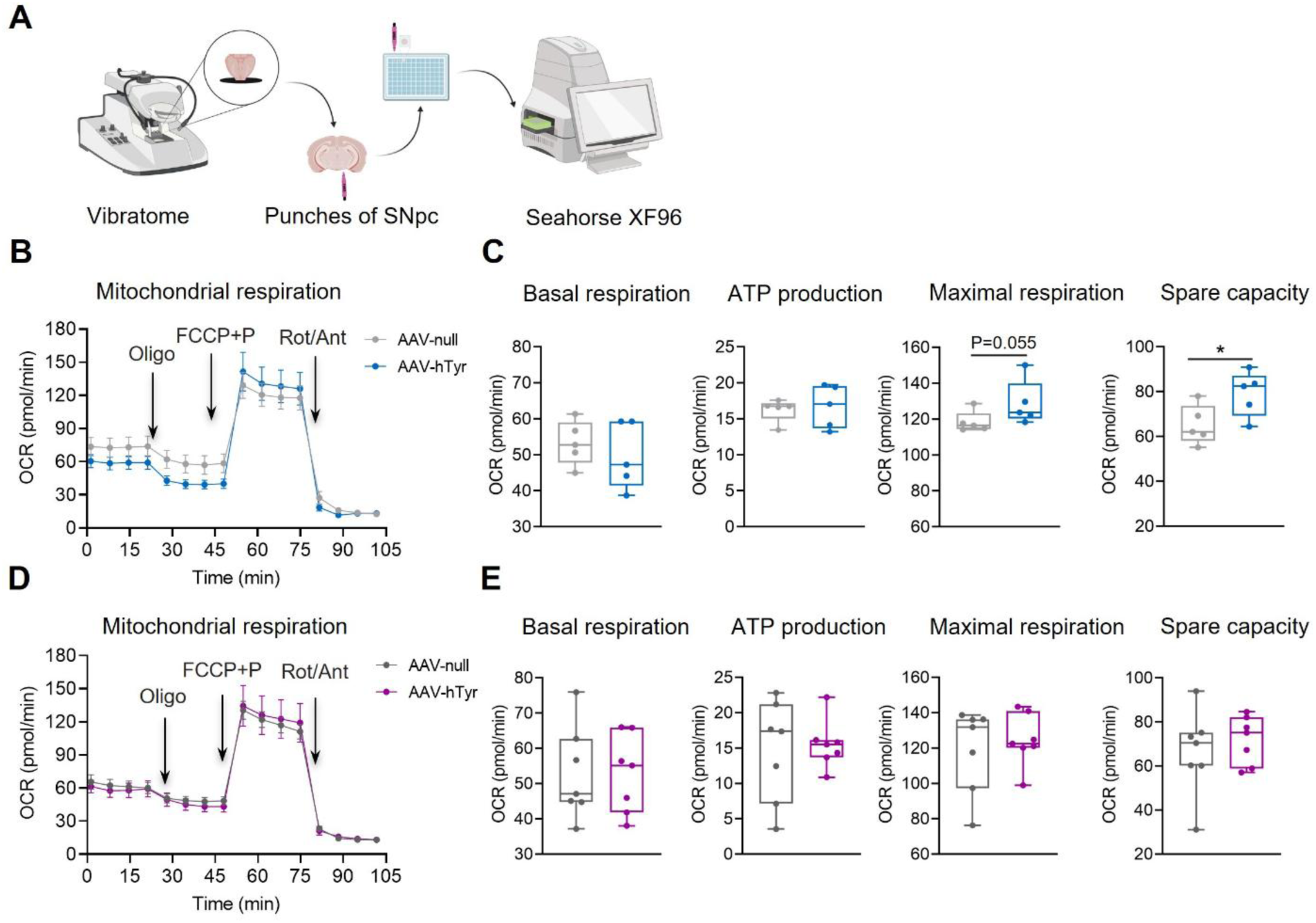
Nigral NM buildup induces sex-specific changes of the mitochondrial functionality in SNpc. **A,** Scheme of the protocol used to evaluate the mitochondrial function on SNpc punches obtained by midbrain rat slices. **B,** Metabolic profile of the Mitochondrial Stress Test showing real-time measurements of the oxygen consumption rate (OCR) after the injection of 1) Oligomycin (Oligo), 2) FCCP pyruvate (FCCP+P), and 3) Rotenone/Antimycin (Rot/Ant) in SNpc punches of male animals. **C,** Mitochondrial parameters obtained from Seahorse mitochondrial stress test in male AAV-hTyr-injected rats (*n* = 5) and AAV-null rats (*n* = 5): basal respiration (P = 0.494, Student’s *t*-test), ATP production (P = 0.690, Mann-Whitney test), maximal respiration (P = 0.055, Mann-Whitney test), and spare respiratory capacity (P = 0.047, Student’s *t*-test). Data are represented as median ± quartiles. *P < 0.05. **D,** Metabolic profile of Mitochondrial Stress Test showing real-time measurements of the OCR after the injection of Oligo, FCCP+P, and Rot/Ant in SNpc punches of female rats. **E,** Mitochondrial parameters obtained from Seahorse mitochondrial stress test in female AAV-hTyr-injected rats (*n* = 7) and AAV-null rats (*n* = 7): basal respiration (P = 0.997, Student’s *t*-test), ATP production (P = 0.766, Student’s *t*-test), maximal respiration (P = 0.620, Student’s *t*-test), spare respiratory capacity (P = 0.527, Student’s *t*-test). Data are expressed as median ± quartiles.

### NM buildup induces sex-specific alterations of nigral DA neurons’ activity

To reveal the impact of NM buildup on functional features of nigral DA neurons, we performed patchclamp recordings in midbrain slices from AAV-hTyr-injected rats and controls 4 weeks post-AAVs injection. First, we analyzed nigral DA neurons’ function in female rats (**Fig. 4A**). NM production was visible in medial SNpc within single DA neurons of AAV-hTyr-injected rats, as dark aggregates close to plasmatic membranes or in the cytosol (**Fig. 4B**); this allowed precise selection of NM-containing neurons to compare with control NM-lacking cells. We found altered passive membrane properties in female NM-producing SNpc DA neurons, displaying reduced membrane capacitance (C_m_), normal membrane resistance (R_m_), and a trend to hyperpolarization (reduced I_hold_ at V_H_ = −60 mV) (**Fig. 4C**). Moreover, NM accumulation deeply perturbed spontaneous firing activity, influencing firing pattern and frequency. As known, nigral DA neurons can exhibit different firing patterns (regular pacemaker, irregular spiking, and bursting) or barely absence of firing (silent neurons) due to influence of multifaceted intrinsic and extrinsic factors. We observed that while in female controls most SNpc DA neurons fired as regular pacemakers and a minority in bursting mode (**Fig. 4D**), NM-producing DA neurons displayed reduced incidence of pacemaker firing neurons and increase of bursting neurons; few cells were irregularly firing or silent (**Fig. 4D**). Analysis of the coefficient of variation of the interspike interval (CV-ISI) confirmed this NM-induced shift in firing patterns, with increased firing irregularity (**Fig. 4E**) and higher bursting activity (**Fig. 4F**) in SNpc DA neurons of AAV-Tyr-injected rats. Additionally, the pacemaker population exhibited a trend toward hyperactivity (**Fig. 4G**).

**Figure 4.**
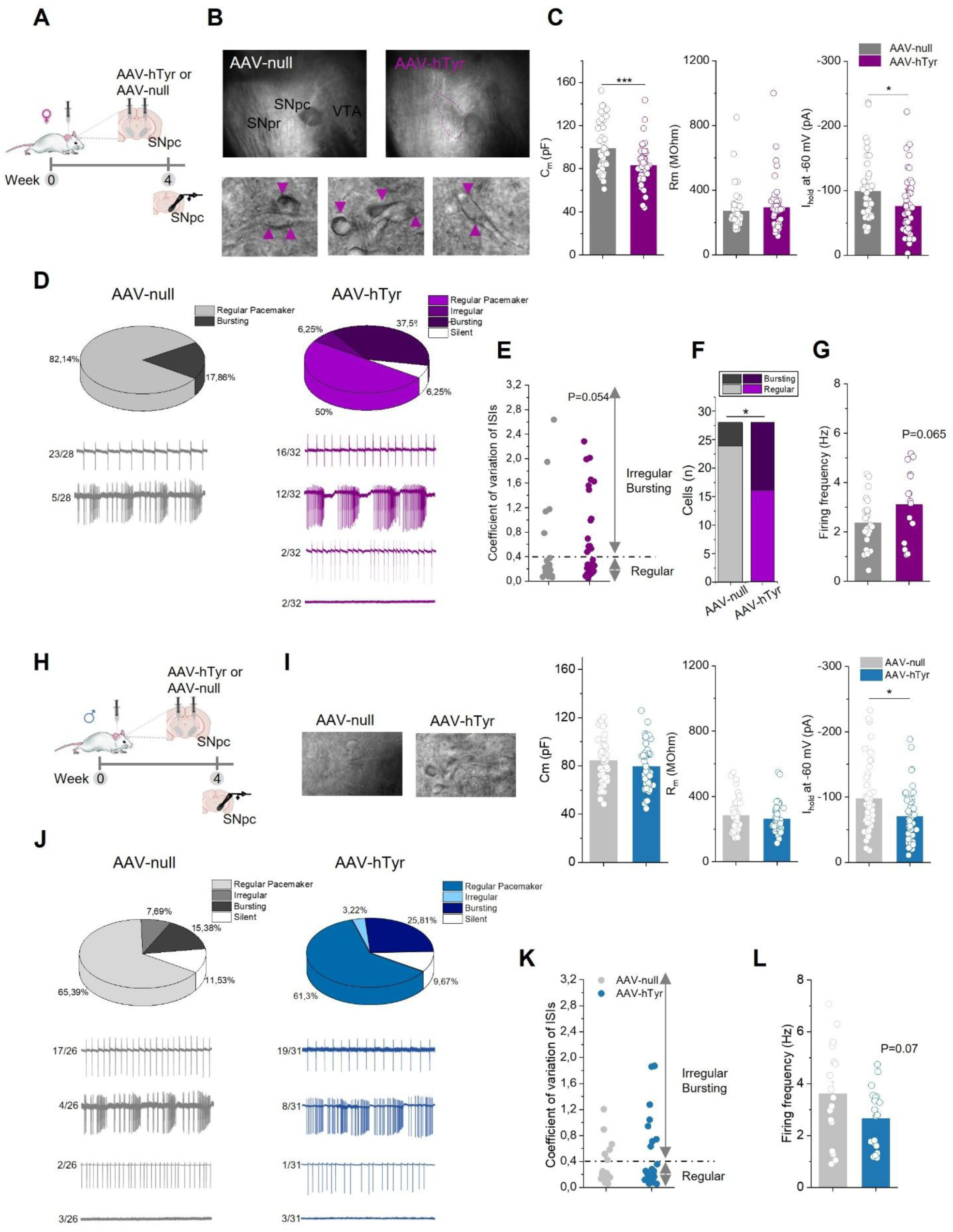
NM induces sex-specific alterations of nigral DA neurons’ activity. **A,** Experimental design used to evaluate NM effects on SNpc DA neurons’ function by *ex vivo* electrophysiology. **B,** Example images of midbrain slices acquired through infrared microscopy showing NM buildup in SNpc (up, 4X magnification) and within nigral DA neurons of AAV-hTyr-injected rats (bottom, 40X magnification). Intracellular NM aggregates are indicated by arrows. **C,** Plots of passive membrane properties of female nigral DA neurons, namely membrane capacitance (C_m_) (left), membrane resistance (R_m_) (middle), and holding current (I_hold_) to V_H_ = −60 mV (right) in AAV-hTyr-injected rats (*n* = 47 cells/3 rats) and AAV-null rats (*n* = 44 cells/3 rats), Mann-Whitney test, C_m_: P = 6.22 × 10^-4^; R_m_: P = 0.387; I_hold_: P = 0.011. **D,** Up, pie diagrams indicating incidence (%) of different firing patterns (regular pacemaker, irregular, bursting, and silent) of female nigral DA neurons; Bottom, Example cell-attached firing traces, with indication of relative prevalence (cells in each firing pattern/total cells). **E,** Histogram of coefficient of variation of interspike intervals (CV-ISIs) showing higher irregularity in female NM-producing DA neurons (AAV-null *n* = 28 and AAV-hTyr *n* = 30, Mann-Whitney test, P = 0.054). **F,** Proportion of regular pacemaker and bursty neurons (χ^2^ test, χ^2^ (1, N=56), P=0.041); **G,** Quantification of firing frequencies of female pacemaker neurons showing a trend to hyperactivity in NM-producing cells (AAV-null *n* = 23 cells/3 rats and AAV-hTyr *n* = 16 cells/3 rats; Student’s *t*-test, P = 0.065). **H,** Experimental design for *ex vivo* electrophysiology of male nigral DA neurons. **I,** Left, Example images of NM-expressing or control SNpc DA neurons during the patch-clamp procedure; Right, Plots of passive membrane properties (C_m_, R_m_, and I_hold_ at V_H_ = −60 mV) of male nigral DA neurons (AAV-null *n* = 49 cells/4 rats and AAV-hTyr *n* = 45 cells/4 rats; MannWhitney test, C_m_: P = 0.215; R_m_: P = 0.591; I_hold_: P = 0.014). **J,** Up, Pie diagrams showing incidence (%) of different firing modes; Bottom, Example firing traces and relative prevalence (cells in each firing mode/total cells). **K,** Plot of CV-ISIs indicating similar temporal spike distribution among male groups (AAV-null *n* = 23 cells and AAV-hTyr *n* = 28 cells, Mann-Whitney test, P = 0.632). **L,** Histogram of firing frequency of male pacemaker neurons showing a trend toward hypoactivity of NM-producing SNpc DA neurons (AAV-null *n* = 17 cells/4 rats; AAV-hTyr *n* = 19 cells/4 rats; Student’s *t*-test, P = 0.073). Data are expressed as mean ± SEM (**C, F, G, I, L**); dots represent individual neurons. *P < 0.05 and ***P < 0.001.

Next, we verified whether NM produces similar functional alterations in male rats (**Fig. 4H**). Male NM-producing DA neurons displayed regular passive membrane properties (normal C_m_ and R_m_) and a trend to hyperpolarization (**Fig. 4I**). NM buildup did not induce substantial alterations in the spontaneous firing pattern of nigral DA neurons, as incidence of regular pacemaker, bursting, irregular, and silent neurons was similar among AAV-hTyr-injected rats and controls (**Fig. 4J**). Coherently, CV-ISIs indicated an identical distribution among regular pacemakers or irregular/bursting cells, with a prevalence of regular pacemakers in both groups (**Fig. 4K**). Remarkably, the pacemaker population displayed a trend toward hypoactivity, oppositely to females (**Fig. 4L**). Collectively, these data prove sex-biased NM roles in shaping nigral DA neurons’ activity.

### Sex-dimorphic effects of NM buildup on intrinsic currents of SNpc DA neurons

To identify mechanisms underlying NM-induced changes of nigral DA neuron activity, we analyzed two intrinsic currents known to shape firing pattern, pacemaker fidelity, and rate: 1) the hyperpolarization-activated current (I_h_) mediated by hyperpolarization-activated cyclic nucleotide-gated (HCN) channels, and 2) the afterhyperpolarization-associated current (I_AHP_), underlying the afterhyperpolarization phase of the action potential^21^. We found that NM buildup did not affect I_h_ in female nigral DA neurons, showing regular I_h_ peak amplitude and density (**Fig. 5A,B**). Differently, NM blunts I_AHP_ area in female SNpc DA neurons (**Fig. 5B**); such decrease was maintained even after normalization over the C_m_ (**Fig. 5C**).

**Figure 5.**
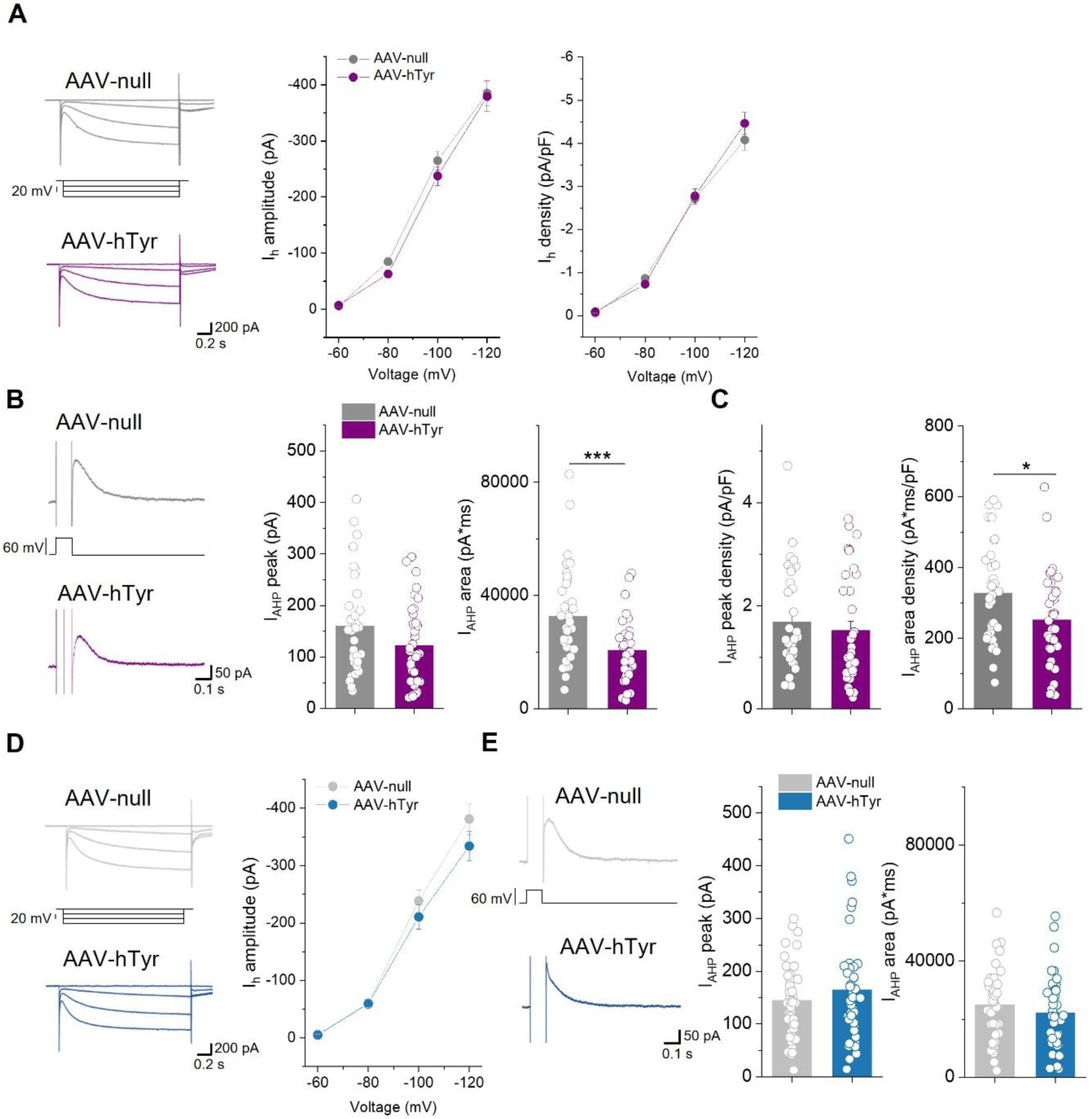
Sex-specific effects of NM buildup on intrinsic currents of SNpc DA neurons. **A,** Example traces of hyperpolarization-activated current (I_h_) (left); quantification of I_h_ amplitude (middle) or I_h_ density (right) of female nigral DA neurons of AAV-null (*n* = 46 cells/3 rats) and AAV-hTyr-injected rats (*n* = 42 cells/3 rats); RM two-way ANOVA, I_h_ amplitude: treatment X voltage P = 0.485. I_h_ density: treatment X voltage P = 0.112. **B,** Representative traces of afterhyperpolarization-associated current (I_AHP_) (left); plots of I_AHP_ peak (middle) and I_AHP_ area (right) in female SNpc DA neurons of AAV-null (*n* = 36 cells/3 rats) and AAV-hTyr rats (*n* = 36 cells/3 rats); Mann-Whitney test; I_AHP_ peak: P = 0.088; I_AHP_ area: P = 8.33 × 10^-4^). **C,** Quantification of I_AHP_ peak and area normalized on C_m_ (I_AHP_ peak density and area density, respectively) (AAV-null *n* = 35 cells/3 rats; AAV-hTyr *n* = 35 cells/3 rats; Mann-Whitney test, I_h_ peak density: P = 0.241; I_h_ area density: P = 0.021. **D,** Example I_h_ traces (left) and quantification of I_h_ amplitude (right) of male nigral DA neurons of AAV-null (*n* = 38 cells/4 rats) and AAV-hTyr rats (*n* = 41 cells/4 rats); RM two-way ANOVA, Treatment X Voltage, P = 0.118). **E,** Representative I_AHP_ traces (left) and plots of I_AHP_ peak (middle) and I_AHP_ area (right) in male nigral DA neurons of AAV-null (*n* = 40 cells/4 rats) and AAV-hTyr (*n* = 38 cells/4 rats); Mann-Whitney test, I_AHP_ peak: P = 0.599; I_AHP_ area: P = 0.292). **A-E,** Data are expressed as mean ± SEM; dots represent individual neurons. *P < 0.05 and ***P < 0.001.

Specular investigations in male rats demonstrated that neither I_h_ (**Fig. 5D**) nor I_AHP_ (**Fig. 5E**) were altered in NM-producing SNpc DA neurons. Thus, NM triggers sex-biased changes in ionic mechanisms that govern neuronal activity. Importantly, I_AHP_ reduction can increase spontaneous firing rate and burstiness, thus explaining the NM-induced shifts of nigral DA neuron activity observed exclusively in females.

### Nigral NM accumulation induces non-motor PD symptoms in male rats

To evaluate whether NM accumulation within SNpc DA neurons contributes to early insurgence of non-motor PD symptoms, we conducted a comprehensive behavioral analysis of AAV-hTyr-injected rats and controls (**Fig. 6A**). Inspection of anxiety domain in the elevated plus maze (EPM) revealed that male NM-producing rats spent less time in the open arms (an index of anxiety) (**Fig. 6B**) and explored less the central area of an open field (OF) (likewise indicative of higher anxiety) compared to controls (**Fig. 6C**). To assess whether nigral NM buildup also promotes social anxiety, male rats were tested in the social avoidance test (SAT). While, as expected, control rats exhibited social preference (that is higher interaction with a social target (ST) over a non-social target (NT)), AAV-hTyr-injected rats displayed social avoidance, spending less time in the interaction zone (IZ) close to ST (**Fig. 6D**). Next, as further scrutiny of anxiety domain, we evaluated compulsive behavior in the marble burying test (MBT), observing a trend, still not significant, to increased number of marbles buried by AAV-hTyr-injected rats (**Fig. 6E**).

**Fig. 6.**
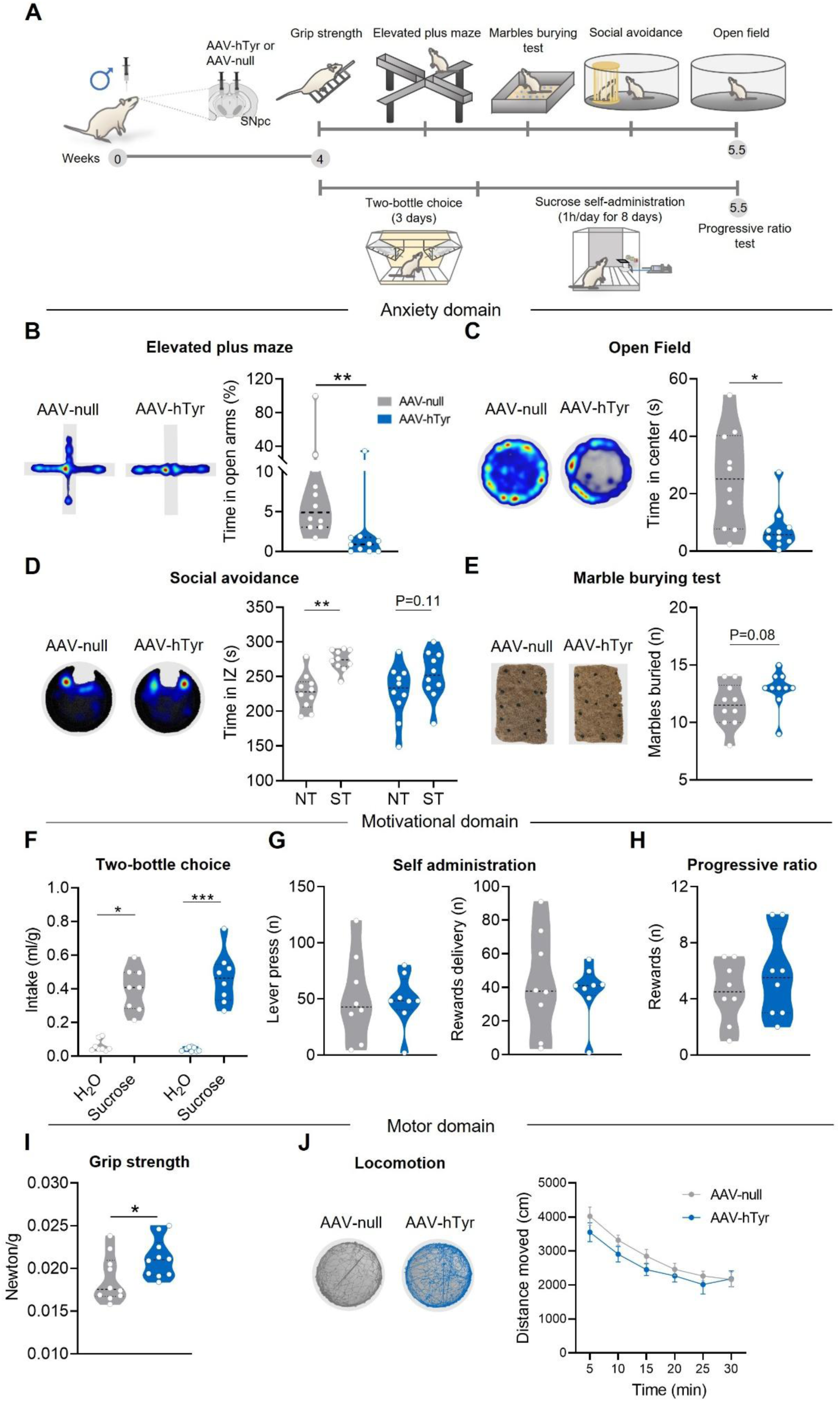
Nigral NM buildup induces non-motor PD symptoms in male rats. **A,** Experimental timeline and design used to evaluate male AAV-hTyr-injected and control rats in different behavioral domains (anxiety, motivational and motor domains). **B,** Heatmaps (left) and quantification (right) of the time spent in the elevated plus maze (EPM) (as % of the total time) showing that AAV-hTyr-injected rats spent less time in the open arms (AAV-null *n* = 10; AAV-hTyr *n* = 10; Mann-Whitney test, P = 0.002). **C,** Heatmaps (left) and quantification (right) of the time spent in the center of an open field (AAV-null *n* = 10 and AAV-hTyr-injected rats *n* = 10, Mann-Whitney test, P = 0.011). **D,** Heatmaps (left) and quantification (right) of the time spent in the interaction zone (IZ) during the social avoidance test; NM-producing rats do not exhibit preference for the social target (ST) over an empty non-social target (NT) (AAV-null *n* = 10; AAV-hTyr *n* = 10; RM two-way ANOVA followed by Sidak’s test, AAV-null: NT *vs* ST P = 0.003, *AAV*-hTyr: NT *vs* ST P = 0.012). **E**, Representative images of marble burying test (MBT) (left) and number of marbles buried (right) (AAV-null *n* = 10 and AAV-hTyr *n* = 10, Student’s *t-*test, P = 0.080). **F,** Quantification of the sucrose intake normalized on the body weight of each animal (ml/g) indicating regular sucrose preference in both groups (AAV-null *n* = 8; AAV-hTyr *n* = 8; Kruskal-Wallis followed by Dunn’s test, H_2_O *vs* Sucrose: AAV-null P = 0.018; AAV hTyr P = 0.001). **G,** Plots of the number of lever press (left) and reward delivery (right) obtained by AAV-null (*n* = 8) and AAV-hTyr-injected rats (*n* = 8). Data are mean values of the last 3 days of the self-administration protocol; comparisons were with Student’s *t*-test (lever press, P = 0.882) or Mann-Whitney test (reward delivery, P = 0.959). **H,** Histogram of the progressive ratio test (AAV-null *n* = 8; AAV-hTyr *n* = 8; Student’s *t*-test, P = 0.413). **I,** Quantification of grip strength normalized on the body weight of each animal (AAV-null *n* = 10; AAV-hTyr *n* = 10; Student’s *t*-test, P = 0.046). **J,** Representative traces (left) and quantification of the distance moved in an open field (right) by AAV-null (*n* = 10) and AAV-hTyr-injected rats (*n* = 10); Mann-Whitney test with Holm-Sidak’s correction: 0-5 min P = 0.538; 5-10 min P = 0.779; 10-15 min P = 0.40; 15-20 min P = 0.895; 20-25 min P = 0.895; 25-30 min P = 0.911). Data are expressed as median ± quartiles **(B-I)** or mean ± SEM **(J)**; dots represent individual animals. *P<0.05, **P<0.01, ***P<0.001.

To assess whether NM buildup also affected the motivational domain, influencing goal-oriented behaviors, rats were subjected to a two-bottle choice test (TBC) for 3 days and to a sucrose self-administration protocol for 8 days (**Fig. 6A**). TBC revealed sucrose preference over water in both groups (**Fig. 6f, Extended Data Fig. 4A)**, also displaying comparable water consumption and body weight gain **(Extended Data Fig. 4B,C)**. This suggests that both groups exhibit preserved hedonic reactions. Outcomes from the self-administration protocol indicated no differences among groups in instrumental responding (number of lever presses and rewards earned) (**Fig. 6G and Extended Data Fig. 4D,E)** nor in motivation, assessed using progressive ratio (**Fig. 6H**). This means that nigral NM buildup at early stages does not affect the motivational domain.

Additionally, we looked over early appearance of motor anomalies in male NM-producing rats. In the grip strength test, AAV-hTyr-injected rats showed greater force than controls (**Fig. 6I),** suggestive of increased rigidity or dysregulation in force’s control. Notably, locomotor activity in an OF was regular, with comparable distance moved in both groups (**Fig.6J**). Altogether, these data demonstrate that NM accumulation within nigral DA neurons promotes early anxiety appearance and slight motor alterations in male rats.

### Female rats are more resilient to nigral NM-induced behavioral alterations

To disclose whether there is sex-biased vulnerability toward NM-induced behavioral alterations, females AAV-hTyr-injected rats and controls were exposed to specular tasks of males (**Fig. 7A**). Female NM-producing rats did not exhibit early signs of anxiety, as proved by equal time spent in EPM open arms (**Fig. 7B**) or in OF center (**Fig. 7C**) with respect of controls. Likewise, female AAV-hTyr-injected rats displayed regular social preference (**Fig. 7D**), and no sign of compulsive behaviors in the MBT (**Fig. 7E**), further confirming absence of anxiety. Additionally, female NM-producing rats exhibited regular sucrose preference (**Fig. 7F, Extended Data Fig. 5A)** and normal performance in sucrose self-administration task, with no differences in lever press, reward delivery (**Fig. 7G, Extended Data Fig. 5B-E),** or progressive ratio test (**Fig. 7H),** thus confirming that nigral NM buildup is influential on goal-oriented behaviors, such as natural rewards seeking. Notably, female NM-producing rats displayed a trend toward increased grip strength (**Fig. 7I**), along with regular locomotion during OF exploration (**Fig. 7J**) compared with controls. Collectively, these findings indicate female rats are less susceptible to early emergence of non-motor PD symptoms during initial stages of nigral NM buildup.

**Fig. 7.**
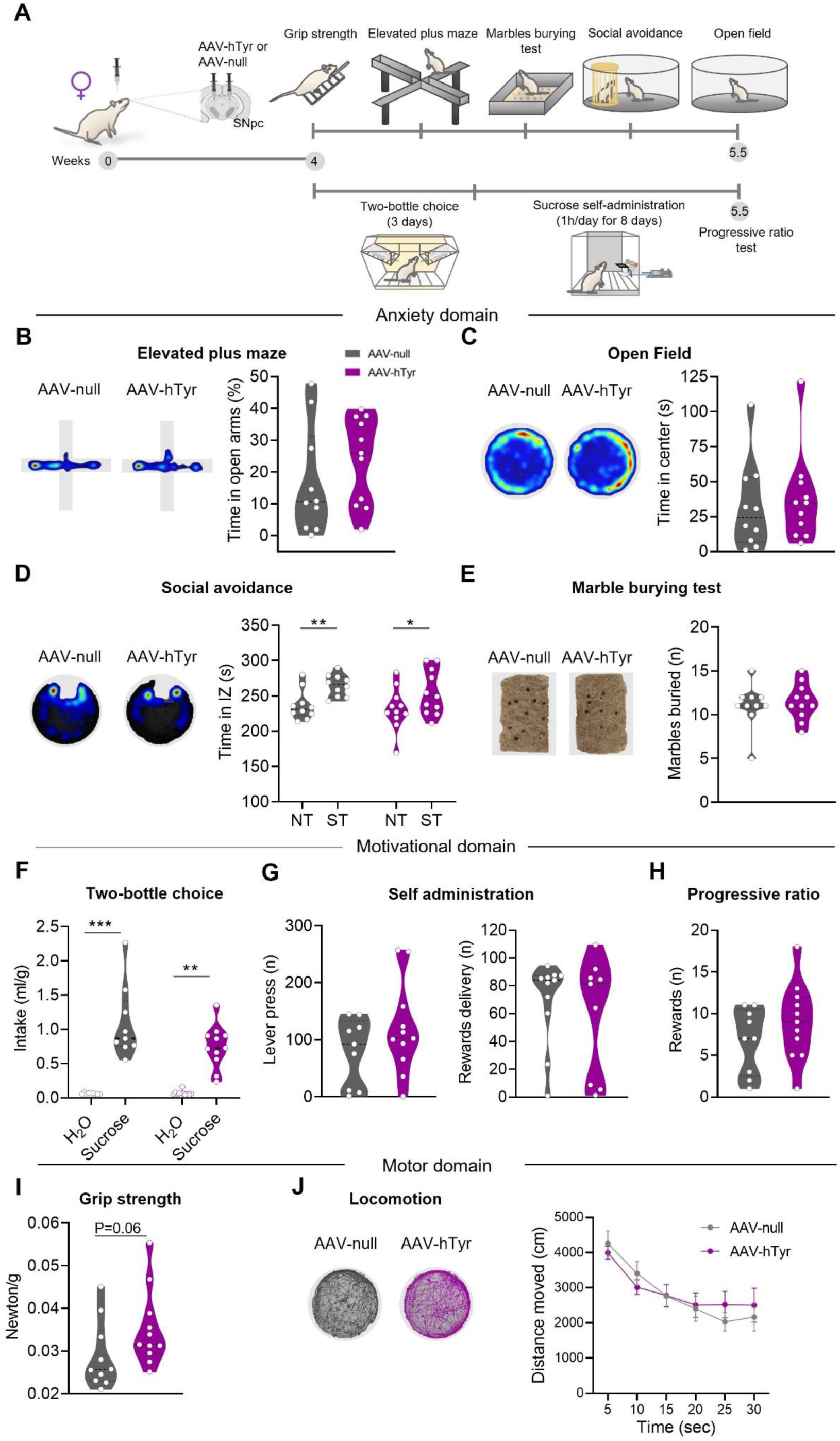
Nigral NM buildup does not induce behavioral alterations in female rats. **A,** Experimental timeline and design used to evaluate female AAV-hTyr-injected and control rats in different behavioral domains (anxiety, motivational and motor domains). **B,** Heatmaps (left) and quantification (right) of the time spent in the elevated plus maze (EPM) (as % of the total time) by AAV-null (*n* = 10) and AAV-hTyr-injected rats (*n* = 11). Student’s t-test, P = 0.297. **C,** Heatmaps (left) and quantification (right) of the time spent in the center of an open field (right) by AAV-null (*n* = 10) and AAV-hTyr-injected rats (*n* = 11); Mann-Whitney test, P = 0.557. **D,** Heatmaps (left) and quantification (right) of the time spent in the interaction zone (IZ) during the social avoidance test, indicating regular preference for the social target (ST) over empty non-social target (NT) in both groups (AAV-null *n* = 10 and AAV-hTyr *n* = 11; RM two-way ANOVA followed by Sidak’s test. AAV-null: NT *vs* ST P = 0.006*; AAV*-hTyr: NT *vs* ST P *=* 0.018). **E**, Example images of marble burying test (MBT) (left) and number of marbles buried (right) by AAV-null (*n* = 10) and AAV-hTyr-injected rats (*n* = 11); Mann-Whitney test (P = 0.845). D, Quantification of the sucrose intake normalized on the body weight of each animal (ml/g) indicating regular sucrose preference in both groups (AAV-null *n* = 9; AAV-hTyr *n* = 11; Kruskal-Wallis followed by Dunn’s test. H_2_O vs Sucrose: AAV-null P = 0.0004; AAV-hTyr P = 0.001). **G,** Plots of the number of lever press (left) and reward delivery (right) obtained by AAV-null (*n* = 9) and AAV-hTyr-injected rats (*n* = 11). Data are mean values of the last 3 days of the self-administration protocol; comparisons were with Student’s *t*-test (lever press, P = 0.236) or Mann-Whitney test (reward delivery, P = 0.710). **H,** Histogram of the progressive ratio test (AAV-null *n* = 9; AAV-hTyr *n* = 11; Student’s *t*-test, P = 0.265). **I,** Quantification of grip strength normalized on the body weight of each animal (AAV-null *n* = 10; AAV-hTyr *n* = 11; Mann-Whitney test, P = 0.06). **J,** Representative traces (left) and quantification (right) of the distance moved in an open field by female AAV-null (*n* = 10) and AAV-hTyr-injected rats (*n* = 11); Mann-Whitney test with Holm-Sidak’s correction (0-5 min P = 0.975; 5-10 min P = 0.823; 10-15 min P = 0.993; 15-20 min P = 0.999; 20-25 min P = 0.913; 25-30 min P = 0.999). Data are expressed as median ± quartiles **(B-I)** or mean ± SEM **(J)**; dots represent individual animals. *P < 0.05, **P < 0.01, ***P < 0.001.

### Sustained NM buildup impairs motor coordination irrespective of sex

To assess whether nigral NM buildup at more advance stages can impair motor function, rats were examined in a rotarod test (measuring motor coordination and learning) 8 weeks after AAV-hTyr injection **(Extended Data Fig. 6A).** Remarkably, prolonged NM overload worsened motor performances on rotarod irrespective of sex **(Extended Data Fig. 6B,C)**. Besides demonstrating that protracted NM accumulation significantly impairs motor function, this evidence suggests that female-biased resilience might be restricted to the initial phases of NM overload, with a later attenuation of sex dimorphism at more advanced stages.

## Discussion

This study shows that NM buildup within SNpc DA neurons triggers multifaceted and sex-specific cellular and behavioral alterations. Specifically, NM affects nigral DA neurons’ activity, mitochondrial functions in SNpc, and fosters non-motor alterations in a sex-dependent manner. Beyond revealing unrealized effects of NM accumulation within nigral DA neurons, our evidence advances the understanding of sex-specific features underlying sex-biased vulnerability in PD.

### NM effects on the nigrostriatal DA circuit

Disclosure of factual NM influence on the nigrostriatal DA system is key to comprehend PD development. While earlier observation that melanized nigral DA neurons preferentially degenerate during PD progression^14^ provided initial clues on NM association to PD, latest evidence envisages multilayered NM effects (protective or detrimental) drawing a more complex picture on NM impact on nigral DA neurons^6,22^. NM at low levels is believed to be neuroprotective, by detoxification of oxidized DA species (an intrinsic process of NM biosynthesis) or metals’ chelation, thus blunting ROS levels and metal-mediated oxidations^22,23^. Otherwise, excessive NM amounts can damage nigral DA neurons, by troubling several cellular processes^6^. Indeed, PD-related dysfunctions in NM-producing rodents and non-human primates (NHP) infer NM contribution to PD development^16,17,18,19,24^. Still, various NM-related aspects, including its impact on neuronal activity or existence of sex-specific factors shaping its roles, need to be clarified. Here, we reveal early sex dimorphic NM effects on nigrostriatal DA system by using a hTyr-based model displaying significant NM levels within nigral DA neurons four weeks post AAV-hTyr injection. Biological sex seems to affect NM biosynthetic rate, as intracellular NM levels are higher in male SNpc DA neurons than in females. In contrast, AAV-hTyr-injected NHP seem to display higher NM levels in female SNpc DA neurons^24^. In our model, nigral NM production induces TH down-regulation irrespective of sex, without significant SNpc DA neurons’ degeneration. A causal link between NM accumulation and TH down-regulation is also supported by evidence showing that pigmentation is associated with fewer TH+ neurons in rodents, NHP and humans^16,17,18,24,25^.

Mitochondrial dysfunction is among the key determinants of PD pathology; impaired mitochondrial respiration in parallel with mitochondrial DNA (mtDNA) deletions has been steadily reported in SNpc in PD patients and models^20,26,27,28,29^. Nevertheless, knowledge about mitochondrial functionality during prodromal PD stages is scarce. We show that NM induces early sex-specific mitochondrial alterations in SNpc, namely increased spare capacity and a trend toward higher maximal respiration in males, but regular functioning in females. This male-specific mitochondrial dysfunction may rely on an adaptation of nigral DA neurons to preserve proper bioenergetic levels to cope with NM burden, as energy-demanding cellular processes (proteasomal and lysosomal degradation) may feasibly trigger to offset NM levels. Notably, similar mitochondrial dysfunctions occur in peripheral blood mon-onucleate cells (PBMCs)^30^, immortalized lymphocytes^31^, and fibroblasts from PD patients with idiopathic or familiar PD forms^32^, indicating commonality amongst peripheral cells and SNpc. Nonetheless, additional in-deep analyses during prodromal and late PD phases require to better disclose mitochondrial alterations developing during PD progression. Assuming that the mitochondrial alterations herein described refer to SNpc DA neurons, input of other nigral cellular types, including astrocytes and microglia, cannot be excluded. Although glial cells may be activated by NM released to the extracellular space from dying DA neurons^33^, NM intracellular buildup in initial stages does not induce neurodegeneration; this narrows the involvement of other cellular types, ascribing mitochondrial dysfunctions to nigral DA neurons, which intrinsically produce NM. Of note, intact mitochondrial function in female neurons, despite significant NM buildup, supports lesser vulnerability compared to males. NM role in fostering mitochondrial dysfunction is reinforced by evidence that pigmented DA neurons of human SNpc show the highest levels of mitochondrial DNA deletions (a vulnerability trait for PD development)^34^ and that NM promotes oxidative stress and impair mitochondrial functions in SH-SY5Y cells, used as proxy of DA neurons^16,35,36^.

### Sex-biased roles for NM in shaping nigral DA neurons’ activity

Adequate comprehension of NM roles in PD pathogenesis cannot overlook the key question on whether (and how) NM buildup influences nigral DA neuron activity. To directly face this unsolved point, we analyzed functional features of NM-producing SNpc DA cells, providing the first evidence of sex-dysmorphic NM impact on nigral DA neuron’s function. Specifically, female NM-accumulating DA neurons display altered passive membrane properties (reduced C_m_), a trend to hyperpolarization, and robust adjustments in the spontaneous firing activity, with a switch in firing patterns from regular pacemaker to bursting. Differently, male NM-producing DA neurons show slighter functional changes, without gross adjustments in the firing pattern, but, oppositely to females, a trend toward hypoactivity.

Regarding NM impact on membrane properties, since C_m_ depends on membrane size and composition, lower values in pigmented female neurons could rely on cell shrinkage and/or altered lipidic bilayer composition or thickness due to NM intercalation. Based on its intrinsic lipidic content, NM can aggregate with lipidic cellular compartments; indeed, NM closeness to DA neurons’ plasma membrane is evident in AAV-hTyr-injected rats. Future studies may gain insight into sex-specific features underlying milder NM influence on C_m_ in male neurons (despite higher NM levels), disclosing whether differences arise from sex dimorphism in membrane composition, favoring NM’s membrane anchoring in females.

NM-producing SNpc DA neurons display a trend toward hyperpolarization. While this could appear at odds with hyperactivity/burstiness of female neurons, a certain level of hyperpolarization needs to allow bursts’ trigger in SNpc DA neurons *ex vivo* slice preparations^37,38^_;_ in this view, mild hyperpolarization stands as a permissive factor to bursts’ onset. Importantly, NM load induces sex-biased opposite effects on spontaneous firing activity of SNpc DA neurons, namely hyperactivity and higher burstiness in females and a trend to hypoactivity in males. To unveil mechanisms underlying such sex-dimorphic NM effects we analyzed two intrinsic currents - I_h_ and I_AHP_ – known to shape firing rate, regularity, and pattern of nigral DA neurons^21^. Regardless of sex, NM does not alter I_h_ in nigral DA neurons. This aligns with regular I_h_ in SNpc DA neurons in PD models based on α-syn accumulation^39,40,41,42^. Otherwise, I_AHP_ is selectively reduced in female NM-producing neurons, while it is unaltered in males. Thus, NM load induces sex-biased I_AHP_ reduction, and this fits with hyperactivity and burstiness of female NM-loaded neurons. Of note, I_AHP_ decrease associated with hyperactivity/burstiness also occur in aged α-syn overexpressing rats^41^, thus denoting shared alterations within PD models. The exact mechanism by which NM affects I_AHP_ remains to be specified; theoretically, NM may inhibit I_AHP_-mediating channels, namely small conductance- and big conductance K^+^ channels (SK and BK) either by a direct binding or by indirect interference with channels’ regulatory mechanisms, i.e., regulatory subunits or intracellular Ca^2+^ fluctuations.

### Sex-dimorphic effects of NM on PD symptomatology

Which is the behavioral phenotype associated with NM buildup within SNpc DA neurons? Do the sex-dimorphic adaptations of nigral DA neurons lie behind sex-biased vulnerability to develop PD symptoms? As known, PD is characterized by a prolonged prodromal period with a high prevalence of neuropsychiatric symptoms, like anxiety, social phobia, depression, and apathy^2^. We show sex-specific vulnerability to develop non-motor PD symptoms during early NM buildup. Increased anxiety is exclusively evidenced in male NM-producing rats. Early appearance of anxiety has been seen in other PD models characterized by blunted DA levels in SNpc projections areas^43,44,45,46^ and structural and functional alterations of SNpc correlate with anxiety in traumatic brain injury^47^. Coherently with females’ resilience to NM-induced anxiety, female *Pink1* KO rats (a genetic PD model) do not display anxiety before ageing^48^. A link between NM and anxiety is further suggested by evidence that nigral load of iron (that is primarily bound to NM) correlates with anxiety in PD patients^49^.

Social anxiety is another common prodromal PD symptom. Nigral NM overload causes social avoidance selectively in males, while females exhibit regular social preference. Such sex-dimorphic social avoidance reflects clinical PD phenotypes, with higher prevalence of social phobia in males^50^. Other evidence advises SNpc DA nucleus’ dysregulation can be instrumental to social deficits, as fewer nigral DA neurons have been reported in a model of autism spectrum disorder^51^, and social avoidance has been correlated with nigral mitochondrial dysfunctions in male mice^52^.

Apathy, anhedonia, and depression can develop concurrently with anxiety in PD patients. Irrespective of sex, NM-producing rats display normal hedonic response and motivation for natural rewards, as proved by regular scores in the TBC and sucrose self-administration protocols. Previous evidence suggests that moderate loss of SNpc DA neurons need to impair goal-oriented behaviors, as reduced sucrose self-administration has been reported in PD models with partial lesions of SNpc DA neurons^53,54^, while sucrose preference is preserved in a α-syn-based PD model displaying slight SNpc DA neurons’ loss^55^. Coherently, initial NM buildup, which does not induce nigral DA neurons’ degeneration, does not affect motivated behaviors.

Nigral NM overload induced motor anomalies overt as increased grip strength of NM-producing rats of both sexes. This aligns with other PD rodent models with unilateral lesions^56,57,58^ and with PD patients, exhibiting higher grip force in early pathological stages^59,60,61^. Abnormal grip force can rely on inability to release an applied force due to dysregulation of basal ganglia circuitry, thus promoting rigidity, a cardinal PD hallmark^62,63^. Notably, nigral NM buildup do not influence basal locomotion at initial stages while impair motor coordination at more advanced phases (∼ 8 weeks) irrespective of sex. Thus, the female-specific resilience appears locked to initial phases of NM overload, with a later attenuation of sex dimorphism.

Male sex is the second most relevant risk factor for PD, beyond aging. Aside to a higher prevalence, PD manifests earlier and with greater DA denervation in men. Sex dimorphism is replicated in PD models, as male rats and primates are more susceptible to toxin-induced SNpc DA neurons’ degeneration than females^64,65,66^. Sex-specific features dictating males’ vulnerability are mostly unknown. Estrogens are commonly considered key factors shaping sex-biased liability to PD, based on their neurotrophic, neuroprotective, and modulatory effects on female nigrostriatal DA circuit^67,68,69,70^ Sex chromosome genes can also underpin sex-biased vulnerability of the nigrostriatal DA circuit during PD development^70,71^. Microarray analyses of single DA neurons from human SNpc sections revealed higher levels of PD-associated genes (α-syn and PINK-1)^72^ and lower expression of genes involved in oxidative phosphorylation and synaptic transmission in males in respect with females^73^. Thus, male SNpc DA neurons own a pattern of gene expression that could foster PD development. The Y-chromosome gene (sex-determining region Y, SRY) has been proposed as a male-specific factor contributing to DA system’s vulnerability. SRY is expressed within a subset of DA neurons of the ventral midbrain of males^74^, wherein influences gene transcription of key enzymes of DA metabolism, such us TH, monoamine oxidase A (MAO_A_), and DA decarboxylase^74,75,76^. SRY is upregulated in 6-OHDA- and rotenone-based PD models, whereas its downregulation is sufficient to mitigate SNpc DA neurons’ degeneration and motor deficits of male PD rodents^77^. Future investigations require to identify key factors accounting for sex-biased NM roles on SNpc DA neurons fostering PD development.

## Conclusions and open issues

This study provides novel insights on NM impact on nigral DA neurons, revealing sex-biased functional adaptations contributing to male-specific susceptibility to develop prodromal PD symptoms during initial NM buildup. Hyperactivity and burstiness of nigral DA neurons emerge as adaptations that females, but not males, are able to trigger during NM buildup. Whether this sex dimorphism relies on female-specific features (naturally lacking in males) or on mechanisms intrinsic to both sexes but impaired in males due to their higher vulnerability to NM, remains to be determined.

Noteworthy, sex-specific changes of SNpc DA neurons’ activity associate with sex-biased behaviors of NM-producing rats. Bursting activity of female SNpc DA neurons could ensure sufficient DA levels to allow DA-dependent functions, explaining why females are more resistant to early manifestation of PD symptoms during initial NM buildup. Otherwise, the trend toward hypofunction of male SNpc DA neurons, in addition to TH down-regulation (affecting DA biosynthetic rate), could reduce DA tone in projection areas exposing males to earlier manifestation of PD symptoms.

Chronic NM production induces behavioural alterations also in female rats, revealing an attenuation of sex-dimorphism at advanced stages of NM buildup. Time-dependent acceleration of pathological signs in females during prolonged NM buildup would reflect clinical evidence in PD patients, as females, despite lower initial incidence, display faster disease progression than males. By reproducing male-specific vulnerability observed in PD patients in early stages as well as females’ pathological recruitment along disease progression, this NM-producing rat model represents a precious tool to explore PD etiology and to probe sex-tailored innovative therapeutic approaches.

Conclusively, beyond corroborating the relationship between NM and PD, this study advances the understanding of the determinants of sex-biased vulnerability observed in PD patients. Increased recognition of sex differences could aid in the development of customized approaches to better manage PD symptomatology.

## Author contributions

**Conceptualization:** A.L., S.L.D, M.M.C, M.V., and N.B.M.

**Methodology:** S.L.D, M.M.C, A.L., D.C., A.F., and M.V.

**Investigation:** S.L.D, M.M.C., A.L., F.C., S.S., V.N., and G.D.

Project administration: **A.L.**

**Supervision:** A.L.

**Funding acquisition:** N.B.M. and M.V.

**Visualization:** A.L, S.L.D, and M.M.C.

**Writing of the manuscript:** A.L. and S.L.D.

All authors revised the manuscript and approved the submitted version.

## Acknowledgements

This research was funded by Aligning Science Across Parkinson’s [ASAP-020505] through the Michael J. Fox Foundation for Parkinson’s Research (MJFF). For open access, the author has applied a CC BY public copyright license to all Author Accepted Manuscripts arising from this submission. The study was also supported by NEXTGENERATIONEU (NGEU) and funded by the Ministry of University and Research (MUR), National Recovery and Resilience Plan (NRRP), project MNESYS (PE0000006) – A Multiscale integrated approach to the study of the nervous system in health and disease (DN. 1553 11.10.2022). Sponsors did not play a role in study design, data collection, analysis and interpretation, or writing of this manuscript.

## Data availability

Source data and key resources used in the current study are available as indicated in the Key Resource table posted at the Zenodo repository at https://zenodo.org/records/15528775.

No code was generated for this study; all data cleaning, preprocessing, analysis, and visualization was performed using OriginPro 2024 (OriginLab), Prism10 (GraphPad), Stereoinvestigator 2019.1.3 (MicroBright-Field), ImageJ (NIH), pClamp 10.3 (Molecular Devices), EthoVision XT17 (Noldus).

Additional request of information should be addressed to the corresponding author Ada Ledonne.

## Methods

### Animals

All procedures were carried out following the guidelines on the ethical use of animals from the Council Directive of the European Communities (2010/63/EU) and were approved by the Italian Ministry of Health (Authorization N°78-2023). Wild-type rats (Sprague-Dawley background; RRID:MGI:5651135) were bred in our facility and housed in a temperature-(23 ± 1 ◦C) and humidity-controlled environment (45 %–60 % relative humidity), with a 12 h light/dark cycle. Male and female rats were subjected to surgical procedures for virus injection at 2 months of age and used for different experiments after 4-8 weeks.

### Stereotaxic virus injection

Rats (250-300 g; 2-month-old) were anesthetized with xylazine (Rompun, 20 mg/ml; 0.5 ml/kg; Bayer) and tiletamine/zolazepam (Zoletil, 100 mg/ml; 0.5 ml/kg; Virbac) and then mounted in a stereotaxic frame (David Kopf Instruments, Tujunga, CA, USA) equipped with a rat adapter. Rats were stereotaxically injected into the SNpc (AP = −4.9 mm, ML=1.8 ± mm, DV = −7 mm) with the viral vectors AAV9-CMV-htyr (AAV-htyr) or AAV9-CMV-null (AAV-null) (2 μl for side, 0.4 μL/min, titer 9.11 X10^12^ gc/mL) produced at the Viral Vector Production Unit of the Autonomous University of Barcelona (UPV-UAB, Spain). After injection, the needle was left in place for an additional 5 min to prevent backflow. Rats were used for different experiments 4-8 weeks post-injections.

### Histopathology of nigrostriatal circuit Brain preparation

Rats were intraperitoneally injected with xylazine (Rompun; 20 mg/ml, 0.5 ml/kg; Bayer) and tiletamine/zolazepam (Zoletil; 100mg/ml, 0.5 ml/kg; Virbac). Rats were transcardially perfused with 1% heparin in 0.1 M sodium phosphate buffer (PB) to remove blood-derived antigens that may interfere with target antigens’ detection, followed by 4% paraformaldehyde (PFA) in phosphate buffer (PB; 0.1 M, pH 7.4). Brain samples were postfixed in 4% paraformaldehyde (PFA) in phosphate buffer (PB; 0.1 M, pH 7.4) at 4 °C. To allow deep perfusion throughout the tissue, brain samples were maintained overnight in 4% PFA. Then samples were rinsed three times in PB solution and put in 30% sucrose solution at 4 °C until sinking for cryoprotection. Next, brains were embedded into Tissue-tek O.C.T. (Optimal Cutting Temperature) compound and frozen in isopentane, cooled into liquid nitrogen, and stored at −80 °C. Coronal sections (30 μm) were cut from the anterior part to the posterior part of the brain with a cryostat (Leica) at −20°C. Slices were stored at 4°C in 0.1 M PB containing 0.02% sodium azide before the staining.

### Immunofluorescence

To verify the AAV-hTyr transduction at 4 weeks post-AAV injection, nine coronal midbrain sections with an interval of five sections (to cover the entire SNpc) were stained by double immunofluorescence with antibodies against tyrosine hydroxylase (TH) and human tyrosinase (hTyr) as follows. First, sections were incubated with a blocking solution made with 5% donkey serum and 0.3% TritonX-100 (Sigma) in a PB solution to prevent the non-specific binding of antibodies. Then, primary antibodies anti-TH antibody (1:800, Novus Cat# NB 300-108, RRID:AB_350436) and anti-hTyr (1:250, Thermo Fisher Scientific Cat# MA5-14177, RRID:AB_10980000), diluted in blocking solution, were incubated together over-weekend at 4°C and then incubated for 2h at room temperature with an adequate Alexa Fluor 488 (1:200, Thermo Fisher Scientific Cat# A-21206, RRID:AB_2535792) and 555 secondary antibodies (1:200, Thermo Fisher Scientific Cat# A-31570, RRID:AB_2536180).

Nuclei were stained with DAPI (4’,6-diamidin-2-fenilindole). The specificity of the immunofluorescence labeling was confirmed by omitting primary antibodies and using normal serum instead (negative control). Confocal laser scanner microscopy (Zeiss, Oberkochen, Germany LSM800) was used to acquire images.

### Immunohistochemistry

For stereological cell count in the SNpc, nine slices, with an interval of five sections, and 12 striatal sections per animal were processed for chromogenic immunohistochemistry. First, the endogenous peroxidase was neutralized with a 0.3% H_2_O_2_ solution in PB. Sections were then incubated with a blocking solution made with 5% normal goat serum and 0.3 % Triton X-100 (Sigma) in a PB solution for one hour. Free-floating sections were incubated with the primary anti-TH antibody (1:800, Novus Cat# NB 300-108, RRID:AB_350436), diluted in PB containing 0.3% Triton X-100 overnight at 4°C. After three rinses, sections were incubated with a biotinylated secondary antibody (Millipore Cat# AP132B, RRID:AB_92488) followed by the incubation with extravidin-peroxidase (1:1000, Sigma-Aldrich Cat# e2886, RRID:AB_2620165), and visualized by Vector Blue substrate Kit (Vector Laboratories Cat# SK-4700, RRID:AB_2314425). Slides were then dehydrated in a series of alcohols (50%, 70%, 95%, 100%), cleared in xylenes, and coverslipped with Entellan mounting medium (Sigma).

### Stereological cell count

The three-dimensional optical fractionator stereological probe was used to estimate the number of TH+, Tyr+, and NM+ neurons in SNpc and VTA. We used the Stereo Investigator System (Micro-Bright-Field, Vermont, USA). An optical microscope (Axio Imager M2, Zeiss, Oberkochen, Germany) equipped with a motorized stage, a MAC 6000 controller (Ludl Electronic Products, Ltd) and a camera are connected to software Stereoinvestigator 2019.1.3 (RRID:SCR_018948; https://www.mbfbi-oscience.com/stereo-investigator). The region of interest was defined by TH staining and according to the rat brain Paxinos and Franklin’s atlas and outlined with a 5x objective^41^. The three-dimensional optical fractionator stereological probe (x, y, z dimension of 50×50×30 μm, with a guard zone of 5 μm along the z axis) was used. Cells were marked with a 100X oil objective.

### Masson-Fontana

Midbrain slices were fixed in 4% PFA in 0.1 M PB overnight at 4 °C and then immersed in 30% sucrose solution. Slices were cut into 5 μm-thick sections by a freezing microtome, and NM granules were stained by the Masson-Fontana Staining Kit (DiaPath #010228). This procedure is based on the ability of NM to chelate metals by reducing silver nitrate to a visible metallic state. Staining was performed by incubating the sections with ammoniac solution for 40 min at 56°C, followed by sodium thiosulphate for 2 min and a final counterstain with Kernechtrot for 7 min. Between each step, samples were rinsed in distilled water. Slides were then dehydrated in a series of alcohols (50%, 70%, 95%, 100%), cleared in xylenes, and coverslipped with Entellan mounting medium (Sigma).

### Intracellular NM quantification

30 µm-thick unstained sections representative of the SNpc from different animals were used for intracellular NM quantification. The visualization of unstained NM brown pigment identified melanized catecholaminergic neurons. Bright-field pictures from unstained sections were taken with an optical microscope (Axio Imager M2, Zeiss, Oberkochen, Germany) coupled to a camera. All SNpc NM+ neurons were analysed by means of optical densitometry using ImageJ software (NIH, USA; RRID:SCR_003070; https://imagej.net/) to quantify the intracellular optical density (OD) of NM, as previously reported^18^. Each cell was manually outlined excluding the nucleus; the integrated density value per cell was measured and corrected for non-specific background by subtracting values obtained from the neuropil in the same images. All quantifications were performed in blind for the different experimental groups.

### Tyrosine hydroxylase fibers’ quantification

Sections were photographed with a light microscope (Axio Imager M2, Zeiss, Oberkochen, Germany). Densitometric analysis (OD) of dorsal striatum and nucleus accumbens (core and shell) TH+ fibers were analyzed using the Java image processing and plugin analysis program in ImageJ (NIH, USA; RRID:SCR_003070; https://imagej.net/). Densitometry values were corrected for non-specific background staining by subtracting densitometric values measured from the nearby cortex. The area distinction was manually outlined according to the rat brain Paxinos and Franklin’s atlas. All quantifications were performed blindly for the experimental groups.

### Analyses of mitochondrial function Tissue punches

Rats were anesthetized with isoflurane, decapitated, and the brain was dissected as previously reported^79^. The brain was placed in ice-cold oxygenated artificial cerebrospinal fluid (aCSF: 120 mM NaCl, 3.5 mM KCl,1.3 CaCl_2_, 1 mM MgCl_2_, 0.4 mM H_2_PO_4_, 5 mM HEPES, 10 mM D-glucose, pH 7.4), saturated with 95% O_2_ and 5% CO_2_ for 30 minutes. Coronal brain sections (180 µm thickness) were obtained with a vibratome (VT1200S, Leica, Nussloch, Germany) in ice-cold aCSF under continuous oxygenation. A reusable biopsy punch (0.35 mm diameter) (World Precision Instrument, FL, USA) was gently pressed on the brain slices to collect punches including the SNpc.

### Seahorse XFe96 analysis

The punches of SNpc were carefully placed in the bottom center of a well of XFe96 Cell Culture Microplate (Agilent Technologies, CA; RRID:SCR_019545; https://www.agilent.com/) filled with 180 µl aCSF with 1 mM sodium pyruvate solution (Gibco™,Grand Island, NY and Scotland, UK). Next, the XFe96 Cell Culture Microplate was placed in a non-CO_2_ incubator at 37°C for 30 minutes. For the calibration, each injection port (A, B, and C) of the XFe96 cartridge was differently loaded with: oligomycin 5 μM (Sigma Aldrich, USA), FCCP (2-[2-[4-(trifluoromethoxy) phenyl] hydrazinylidene]-propanedinitrile) 1 μM (Sigma Aldrich, USA) + Pyruvate 0.75 μM (Gibco™,Grand Island, NY and Scotland, UK), or rotenone 1 µM (Sigma Aldrich USA) /antimycin A 1 µM (Sigma Aldrich, USA). As soon as the calibration was completed, the plate was extracted and the XFe96 Cell Culture Microplate containing tissue punches was inserted to start the assay of mitochondrial functionality. The test consisted of 4 cycles of measurements (each consisting of: 2 minutes of mix, 1 minute of wait, and 3 minutes of measurement) to assess the oxygen consumption rate (OCR) before (baseline) and after injection of different drugs.

### Electrophysiology Slice preparation

Acute midbrain slices containing the SNpc were used for electrophysiological patch-clamp recordings of nigral DA neurons of AAV-null and AAV-hTyr-injected rats. Midbrain slices were obtained following published procedures^41^ with minor changes. Briefly, rats were anesthetized with isoflurane and decapitated. The brain was rapidly removed from the skull, and a tissue block containing the midbrain was immersed in a low-sodium N-methyl-D-glucamine (NMDG)-based artificial cerebrospinal fluid (aCSF) at 2-4 °C. The NMDG-based solution contained (in mM): 92 NMDG, 2.5 KCl, 1.25 NaH_2_PO_4_, 30 NaHCO_3_, 20 HEPES, 25 glucose, 2 thiourea, 5 Na-ascorbate, 3 Na-pyruvate, 0.5 CaCl_2_, and 10 MgSO_4_, saturated with 95% O_2_–5% CO_2_ (pH 7.3). Horizontal slices (250 μm) were cut using a vibratome (Leica VT1200S, Leica Microsystems, Wetzlar, Germany). Slices were maintained in NMDG-based aCSF at 33.0 ± 0.5°C for 15 min before adding increasing volumes of a sodium-spike solution (2M NaCl in NMDG-based aCSF) every 5 min for 40 min, then were transferred in a HEPES-based aCSF for long-term storage, containing (in mM): 92 NaCl, 2.5 KCl, 1.25 NaH_2_PO_4_, 30 NaHCO_3_, 20 HEPES, 25 glucose, 2 thiourea, 5 Na-ascorbate, 3 Na-pyruvate, 2 CaCl_2_, and 2 MgSO_4_ (pH 7.3). After 1h of recovery, slices were transferred in the recording chamber and perfused at 2.5–3.0 mL/min with aCSF (33.0 ± 0.5°C) containing (in mM): NaCl 126, NaHCO_3_ 24, glucose 10, KCl 2.5, CaCl_2_ 2.4, NaH_2_PO_4_ 1.2 and MgCl_2_ 1.2 (95% O_2_–5% CO_2_, pH 7.3).

### Electrophysiological recordings

Patch-clamp recordings from SNpc DA neurons were performed with setups equipped with upright microscopes (Nikon Eclipse FN1, RRID:SCR_014995; BX51WI Olympus, RRID:SCR_015801) and infrared video-cameras (CoolSNAP Photometrics, and Evolve Photometrics, Arizona). Identification of SNpc DA neurons was done based on their morphology (fusiform, tightly packed, and medium to large-sized cell bodies (diameter ≥ 20 μm)), localization (in the medial SNpc, close to the medial terminal nucleus of the accessory optic tract, MT), and their typical electrophysiological features, as slow spontaneous firing (1–8 Hz) in cell-attached configuration, large action potential (> 2ms) with a prominent afterhyperpolarization, or the presence of a prominent hyperpolarization-activated inward current (I_h_) in response to hyperpolarizing voltage steps^41,79,80^.

Cell-attached and whole-cell patch-clamp recordings were performed with glass borosilicate pipettes (4–7 MΩ) pulled with a PP-83 Narishige puller and filled with a solution containing (in mM): 125 K-gluconate, 10 KCl, 10 HEPES, 2 MgCl_2_, 4 ATP-Mg_2_, 0.3 GTP-Na_3_, 0.75 EGTA, 0.1 CaCl_2_, 10 Phos-phocreatine-Na_2_ (pH 7.3 with 2 M KOH). Recordings were made with a Multiclamp 700B amplifier (Molecular Devices, USA; RRID:SCR_018455) using Clampex 10.3 software (pClamp, Molecular Devices, USA; RRID:SCR_011323, http://www.moleculardevices.com/products/software/pclamp.html) and the A/D converter Digidata 1440A (Molecular Devices, USA; RRID:SCR_021038) connected to a computer.

The spontaneous firing activity was recorded in cell-attached configuration. The mean firing frequency and coefficient of variation of the interspike intervals (CV-ISIs) were measured in at least 2-3 min of recordings. Membrane resistance (R_m_) and capacitance (C_m_) were measured with the “membrane test” protocol (Clampex 10.3) consisting of a 5 mV hyperpolarizing step from −60 mV, at 33.3 Hz by averaging 100 consecutive measurements to calculate final values. R_m_ and C_m_ were measured within 1 min after membrane rupture. Membrane access resistance was repeatedly monitored and recordings in which it exceeded 25 MΩ were discarded. No liquid junction potential correction was applied.

I_h_ was elicited by a voltage-clamp protocol of four hyperpolarizing voltage steps (−60/-120 mV, 20 mV increment, for 1 sec, V_H_= −60 mV) applied to SNpc DA neurons. I_h_ amplitude was the difference between instantaneous and steady-state current at each voltage step. The afterhyperpolarization-associated current (I_AHP_) was induced by applying a depolarizing voltage step (from V_H_ = −60 mV to 0 mV, 100 ms) and measured as I_AHP_ peak and area under the outward current which follows the depolarization.

### Behavior

Behavioral analyses was conducted following the new Research Domain Criteria initiative of National Institute of Mental Health (NIMH RDoC, https://www.nimh.nih.gov/research/research-funded-by-nimh/rdoc/index.shtml). Behavioural tests were conducted between 9:00 am and 16:00 pm after 1 hour of room acclimation.

### Grip strength

The test was conducted using the hanging grid as previously described^81^. The average value obtained from three trials was normalized on animal weight to determine the final score.

### Marble burying test

Marble burying test (MBT) was conducted as previously reported^78^, with minor modifications. Briefly, rats were placed individually in a novel cage (43 cm X 26 cm X 20 cm) containing 5 cm of regular bedding and fifteen marbles (15 mm diameter) and left undisturbed for 20 min. The number of marbles buried was estimated by considering buried a marble covered ≥ 2/3 with bedding. Marbles were cleaned with 50% ethanol before each session.

### Elevated plus maze

Elevated plus maze (EPM) was carried out as previously described^82^. The apparatus consisted of 2 closed arms and 2 open arms (each 51 cm × 11 cm). The percentage of entries in the open arms (open entries / open + closed × 100) and the percentage of time spent in the open arms (time in open / open + closed × 100) during 5 min of exploration were collected and analysed by the automated video-tracking system EthoVision XT17 (Noldus, The Netherlands; RRID:SCR_000441; https://www.noldus.com/ethovision).

### Social avoidance test

The social avoidance test (SAT) was conducted following published procedures^83^, with minor changes. A rat was placed in a circular open field arena (88 cm diameter) containing a protective cage (13 cm × 13 cm) placed on a side of the arena, defined as interaction zone (IZ). The test consists of two sessions of exploration of 5 min each. The first session with an empty cage (no target, NT) and the second session with a social target (ST) represented by a novel rat matched for age and sex. The total time spent in the IZ was recorded and calculated by the fully automated video tracking system EthoVision XT17 (Noldus, The Netherlands; RRID:SCR_000441; https://www.noldus.com/ethovision).

### Open field

Locomotor activity was assessed in an open field (OF) arena (diameter 100 cm), by evaluating total distance moved and time spent in the arena center during 30 min of exploration.

### Sucrose self-administration

#### Self-administration apparatus

Rats were trained in self-administration chambers located inside sound-attenuating cubicles, fitted with an electric fan and controlled by a custom-made system. Each self-administration chamber was equipped with a stainless-steel grid floor and two operant panels placed on the left and right walls. The right panel was equipped with the palatable solution-paired active (retractable) lever. Responses on this lever activated the infusion pump and the three-light cue located above the lever. The palatable solution was delivered to a receptacle located near the solution-paired lever, connected with a silicon tubing to a syringe that contained the palatable solution (see Fig. 6a).

#### Two-bottle choice

The training procedure was as previously described^53^. Two-bottle choice (TBC) started 3 days before self-administration procedures. The first day rats were individually housed with two bottles containing water (habituation). After habituation, one bottle was replaced with another one containing a sucrose solution at 2.5%. The intake of each animal was measured for two consecutive days and then normalized for the body weight.

#### Self-administration training

Rats were trained to self-administer the sucrose solution 1 hour/day for 8 days (first and second day were trained for 3h) at fixed ratio 1 (FR1) with no timeout^53^. The training sessions started with the illumination of the house light and the insertion of the sucrose solutionpaired lever; responses on this lever resulted in the delivery of the sucrose solution (0.2 mL for 4 s) paired with the illumination of the three-light cue (for 20 s).

#### Progressive ratio test

After the self-administration training, rats were tested for progressive ratio (PR), an index of motivation. In this procedure, the number of lever presses (ratio value) required for drug delivery was incremented within sessions and different days following this progression: 1, 2, 4, 6, 9, 12, 15, 20, 25, 32, 40, 50, 62, 77, 95, 118, 145, 178, 219, 268, 328, 402, 492, 603, etc.^84^. The completion of each ratio requirement resulted in the delivery of the sucrose solution (0.2 mL). Sessions lasted 3h; then, the levers were retracted. Rats were excluded when 1 hour passed from the last reward earned.

#### Rotarod test

The test was performed as previously described^55^ with minor modifications. Animals were accustomed to being placed on the rod rotating at low speed (4 rpm) for at least 30 seconds before the first session of the test. The test consisted of three sessions performed on three consecutive days; each session consisted of three trials with an accelerating rod from 4 to 40 rpm in 300 s, and at least 5 min of rest between trials. The time in which rats fell from the rod was recorded for each trial and the average of the three daily trials was used as final value.

## Statistical analysis

Statistical analyses were conducted using OriginPro 2024 (OriginLab; RRID:SCR_014212; http://www.originlab.com/) and Prism10 (GraphPad; RRID:SCR_002798; http://www.graphpad.com/). The normality of data sets was evaluated with Shapiro-Wilk test and/or D’Agostino & Pearson test. For normally distributed data, comparisons were performed with two-tailed Student’s t-test (simple or with Welch’s correction), repetitive measures two-way ANOVA, as appropriate. When normality was violated, data were analyzed with Mann-Whitney test, Chi-Square test, Kruskal-Wallis ANOVA followed by post hoc Dunn’s multiple-comparison, Friedman ANOVA followed by post hoc Wilcoxon signed-rank test (for within paired comparisons) and Kolmogorov– Smirnov test, or Mann-Whitney test with Holm-Sidak multiple-comparison method (for between unpaired comparisons).

## Extended Data Figures

**Extended Data Fig. 1.**
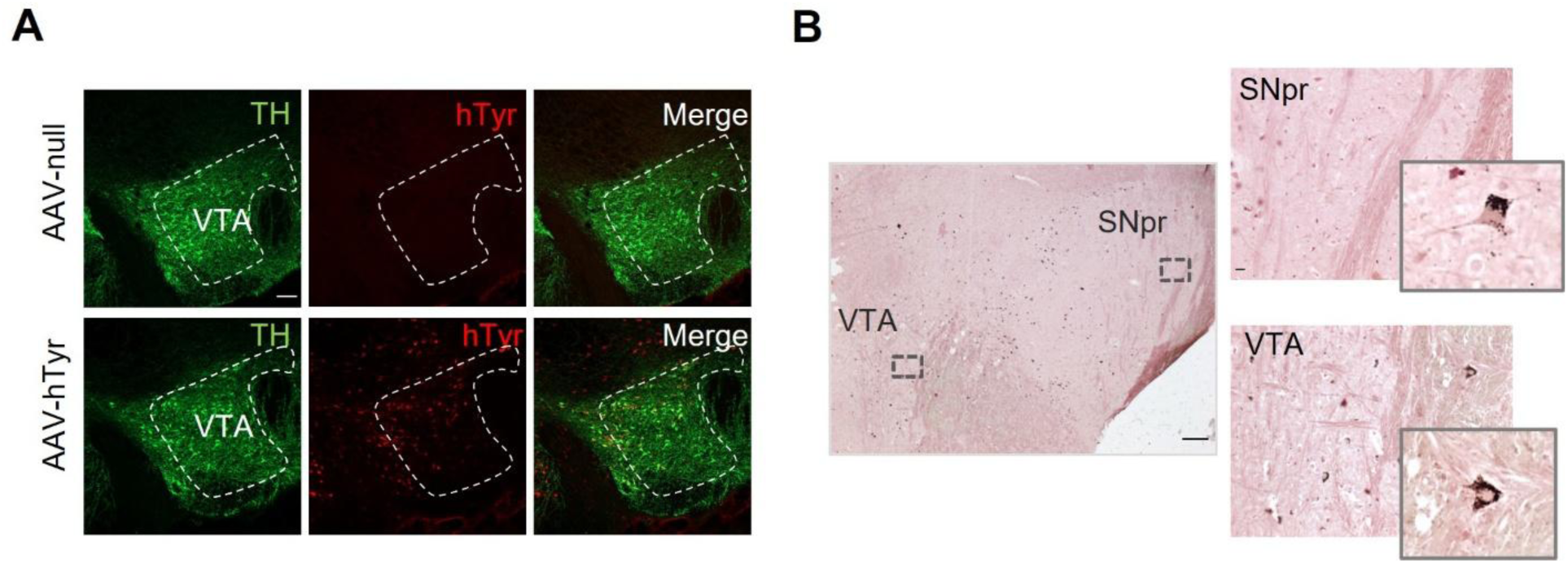
AAV-induced hTyr expression and NM accumulation in VTA and SNpr **A,** Representative double-labeled confocal images of coronal VTA sections immunostained for TH (green) and hTyr (red) (scale bar: 100 μm). **B,** Representative micrographs of Masson-Fontana staining in midbrain sections from an AAV-hTyr-injected rat, showing NM accumulation (in dark brown) in VTA and SNpr four weeks post-AAV-hTyr injection (scale bar: 200 μm, 20 μm, and 10 μm).

**Extended Data Fig. 2.**
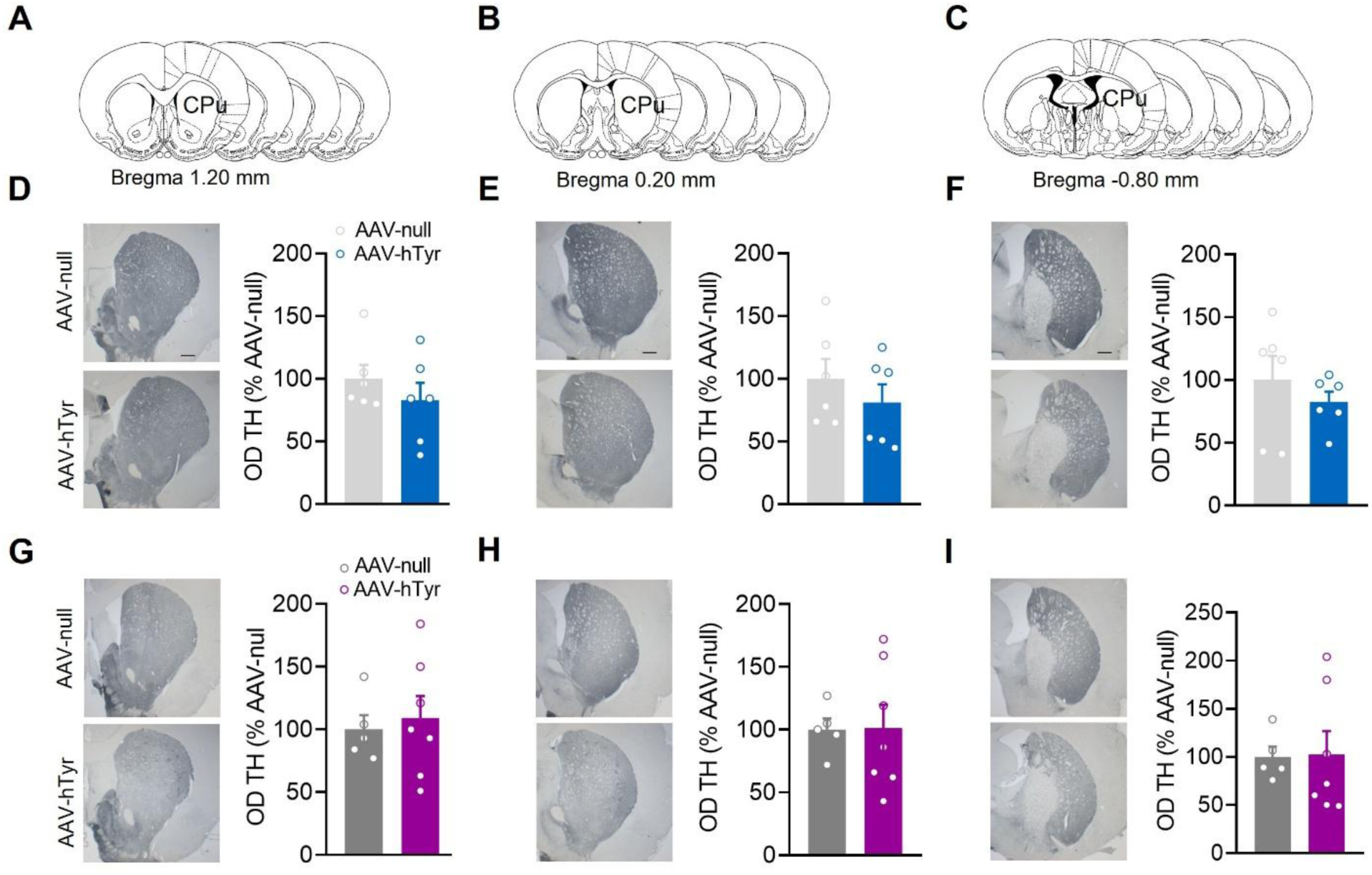
Nigral NM buildup does not impair dST dopaminergic fibers. **A-C,** Qualitative rat brain atlas showing analysed bregma ranges of coronal serial sections of dorsal striatum (dST) for quantitative histological analysis. **D-F,** Representative TH-immunolabeled striatal coronal sections (left, scale bar: 500 μm), and quantification (right) of TH fibers optical density (as % of AAV-null) in male rats (AAV-null *n* = 6; AAV-hTyr *n* = 6; d: Mann-Whitney test, P = 0.562; e: Student’s *t*-test, P = 0.398; f: Student’s *t*-test, P = 0.417. **G-I,** Representative TH-immunolabeled striatal coronal sections (left, scale bar: 500 μm), and quantification (right) of TH fibers optical density (as % of AAV-null) in female AAV-null (*n* = 5) and AAV-hTyr-injected rats (*n* = 7); g: Student’s *t*-test, P = 0.712; h: Student’s *t*-test, P = 0.967; i: Student’s *t*-test, P = 0.929. Data are expressed as mean ± SEM **(D-I)**; dots represent individual animals.

**Extended Data Fig. 3.**
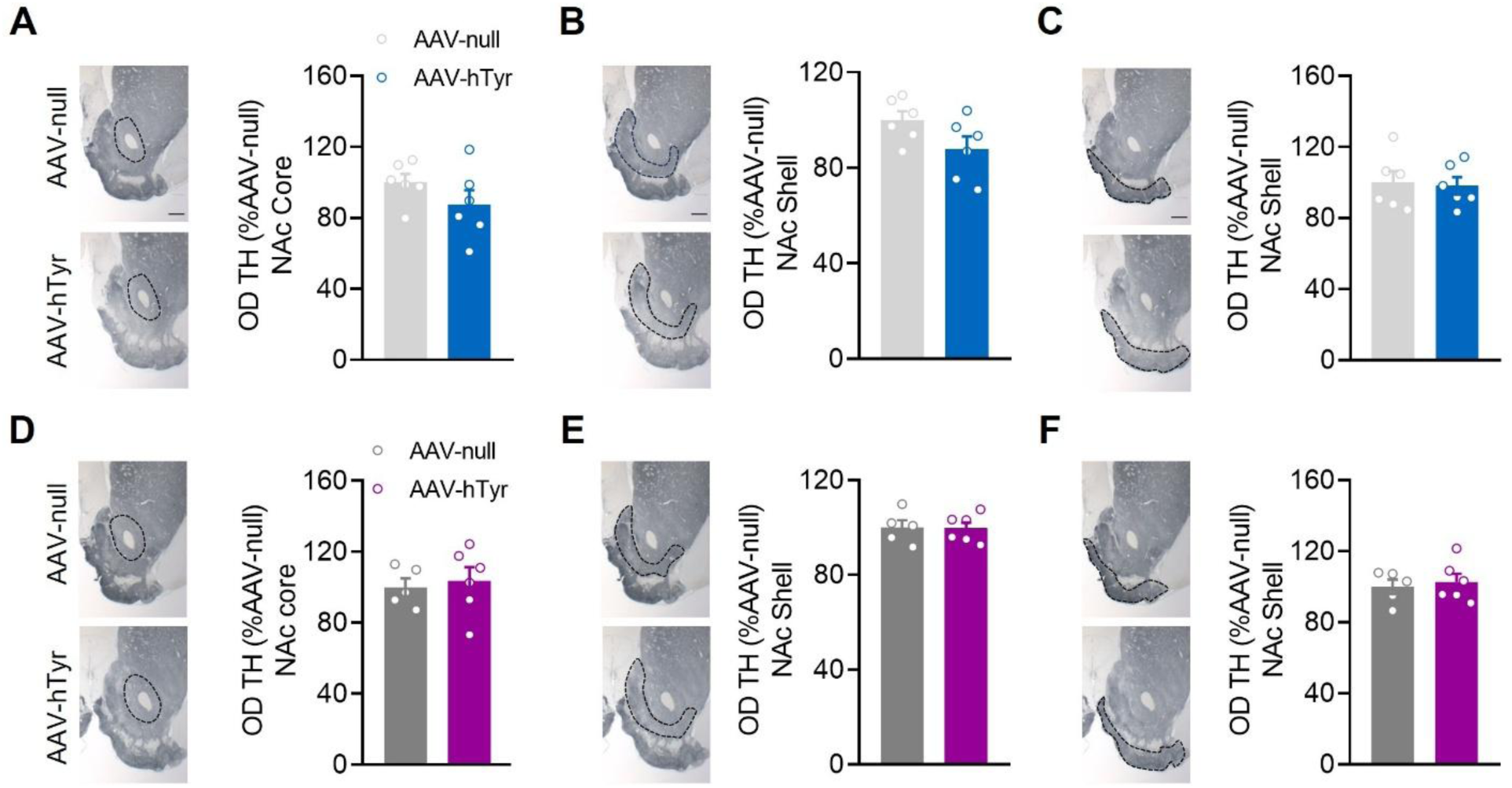
Nigral NM buildup does not impair NAc dopaminergic fibers. **A-C,** Representative TH-immunolabeled coronal sections of NAc Core and NAc Shell (left, scale bar: 500μm), and quantification (right) of TH fibers optical density (as % of AAV-null) in male rats (AAV-null *n* = 6; AAV-hTyr *n* = 6; a: Student’s *t*-test, P = 0.211; b: Student’s *t*-test, P = 0.087; c: Student’s *t*-test, P = 0.818). **D-F,** Representative TH-immunolabeled striatal coronal sections (left, scale bar: 500 μm) and quantification (right) of TH fibers optical density (as % of AAV-null) in female rats (AAV-null *n* = 5; AAV-hTyr *n* = 6; d: Student’s *t*-test, P = 0.708; e: Student’s *t*-test, P = 0.920; f: Student’s *t*-test, P = 0.685). Data are expressed as mean ± SEM **(A-F)**; dots represent individual animals.

**Extended Data Fig. 4.**
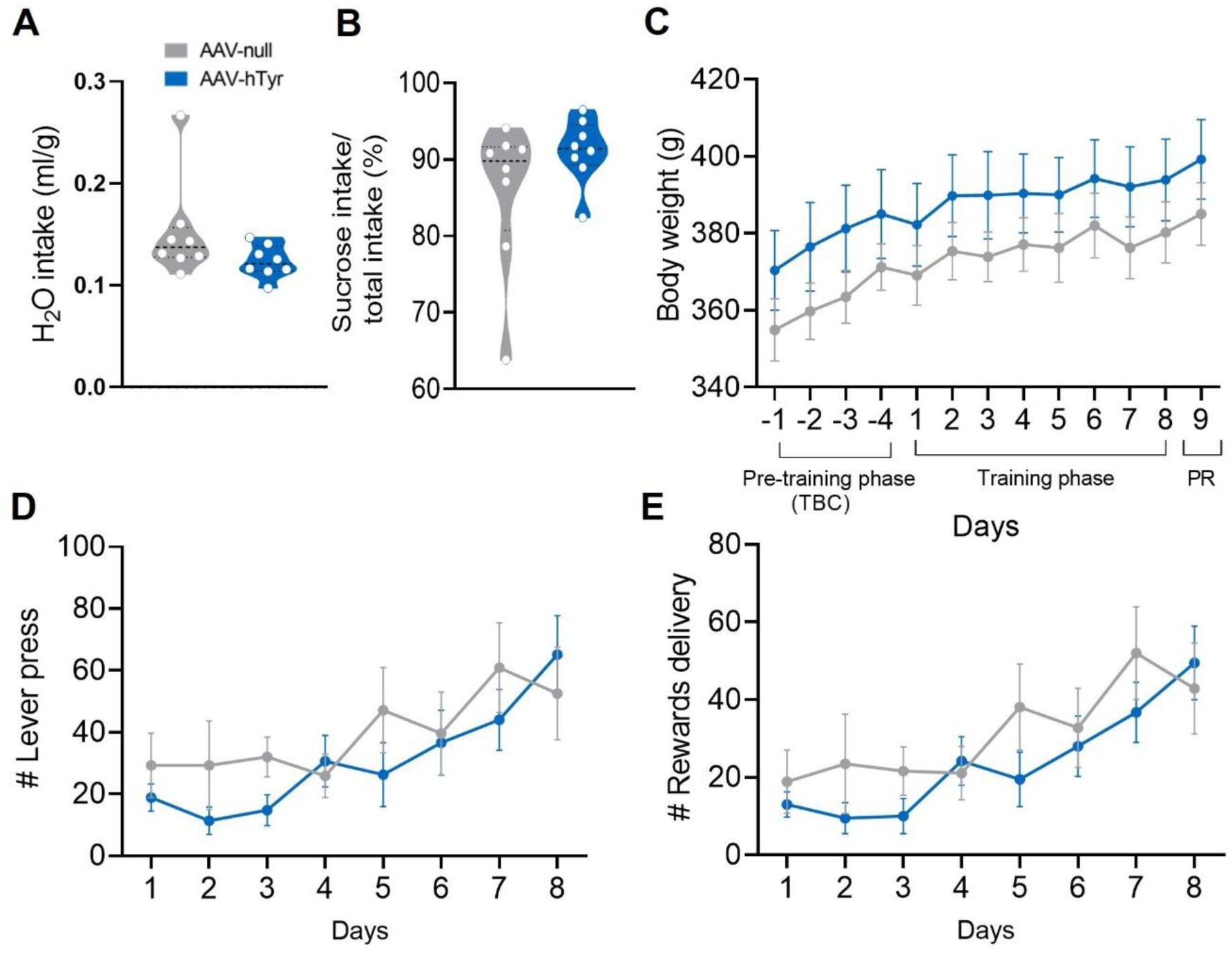
Additional data related to the sucrose self-administration in male rats presented in Fig. 6. **A**, Quantification of water intake normalized on body weight of each animal (ml/g) (AAV-null *n* = 8; AAV-hTyr *n* = 8; Mann-Whitney test, P = 0.130). **B,** Quantification of sucrose intake (as % of total intake) of AAV-null (*n* = 8) and AAV-hTyr-injected rats (*n* = 8); Mann-Whitney test (P = 0.234). **A-B)** Data are median ± quartiles and dots represent individual animals. **C**, Plot of mean body weight of AAV-null (*n* = 8) and AAV-hTyr-injected rats (*n* = 8) during different experimental phases: two-bottle choice (TBC, days −1 to −4), training phase (days 1 to 8), and progressive ratio (PR) (day 9); RM two-way ANOVA followed by Sidak’s test, days: −1 (P = 0.979), −2 (P = 0.973), −3 (P = 0.949), −4 (P = 0.992), 1 (P = 0.995), 2 (P = 0.988), 3 (P = 0.974), 4 (P = 0.991), 5 (P = 0.992), 6 (P = 0.997), 7 (P = 0.975), 8 (P = 0.994) and 9 (P = 0.990). **D**, Chart of mean number of lever press across days (AAV-null *n* = 8; AAV-hTyr *n* = 8; Mann-Whitney test with Holm-Sidak’s correction, days: 1 (P = 0.963), 2 (P = 0.871), 3 (P = 0.163), 4 (P = 0.967), 5 (P = 0.642), 6 (P = 0.967), 7 (P = 0.910), and 8 (P = 0.910). **E**, Plot of mean number of rewards delivery across days (AAV-null *n* = 8; AAV-hTyr *n* = 8; Mann-Whitney test with Holm-Sidak’s correction; days: 1 (P = 0.989), 2 (P = 0.989), 3 (P = 0.718), 4 (P = 0.987), 5 (P = 0.761), 6 (P = 0.989), 7 (P = 0.872) and 8 (P = 0.985). **C-E)** Data are expressed as mean ± SEM. *P < 0.05, **P < 0.01.

**Extended Data Fig. 5.**
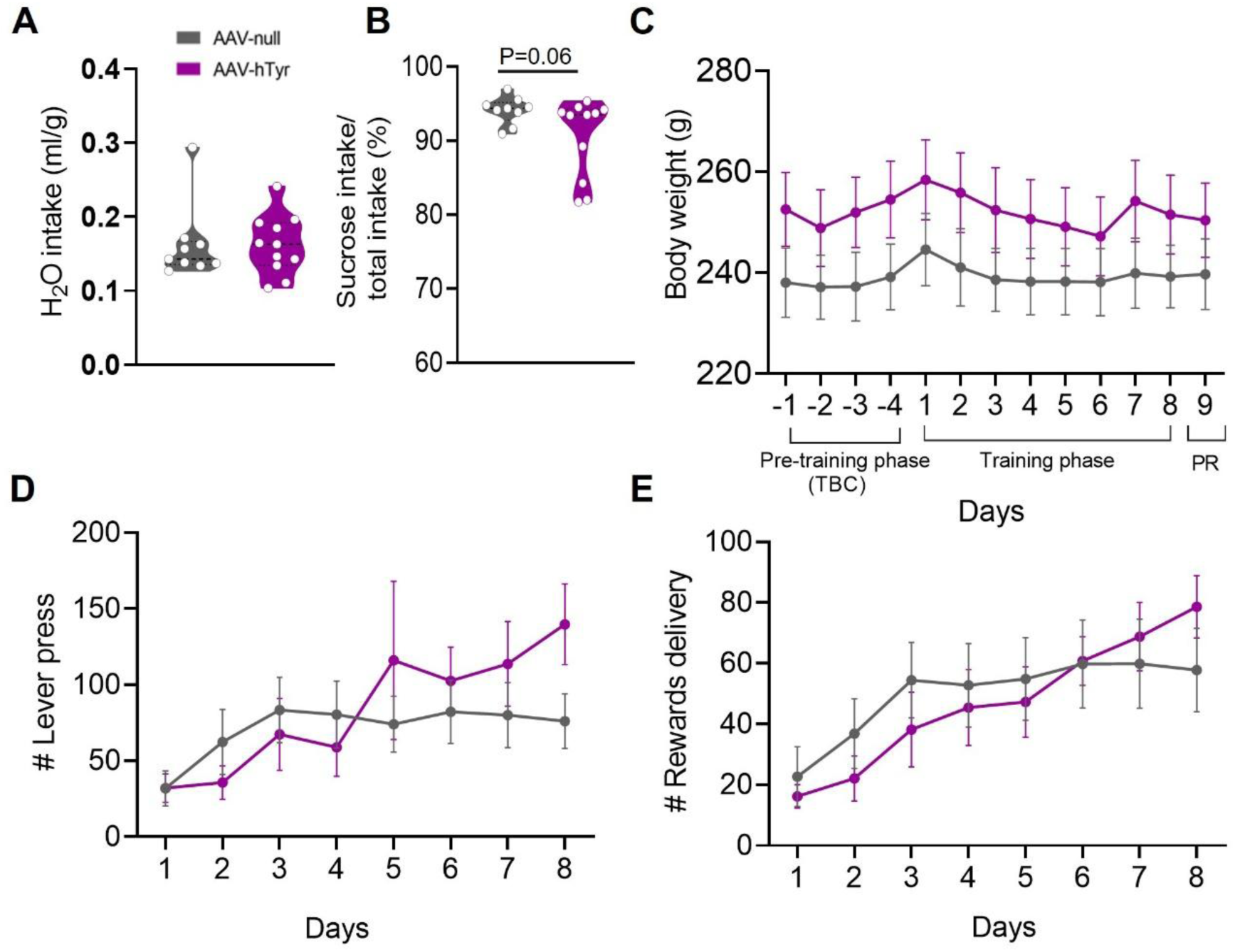
Additional data related to the sucrose self-administration in female rats presented in Fig. 7. **A**, Quantification of water intake normalized on body weight of each animal (ml/g) (AAV-null *n* = 9; AAV-hTyr *n* = 11; Mann-Whitney test, P = 0.564). **B,** Quantification of sucrose intake (as % of total intake) of AAV-null (*n* = 9) and AAV-hTyr-injected rats (*n* = 11); Mann-Whitney test, P = 0.067. **(a,b)** Data are median ± quartiles and dots represent individual animals. **C**, Plot of mean body weight of AAV-null (*n* = 9) and AAV-hTyr-injected rats (*n* = 11) during different experimental phases: two-bottle choice (TBC, days −1 to −4), training phase (days 1 to 8), and progressive ratio (PR) (day 9). Mann-Whitney test with Holm-Sidak’s correction, days: −1 (P = 0.985), −2 (P = 0.985), −3 (P = 0.977), −4 (P = 0.982), 1 (P = 0.985), 2 (P = 0.985), 3 (P = 0.985), 4 (P = 0.985), 5 (P = 0.985), 6 (P = 0.985), 7 (P = 0.985), 8 (P = 0.985), and 9 (P = 0.985). **D**, Chart of mean number of lever press across days (AAV-null *n* = 9; AAV-hTyr *n* = 11; Mann-Whitney test with Holm-Sidak’s correction; days: 1 (P = 0.999), 2 (P = 0.999), 3 (P = 0.998), 4 (P = 0.999), 5 (P = 0.998), 6 (P = 0.999), 7 (P = 0.983), and 8 (P = 0.672). **E**, Plot of mean number of rewards delivery across days (AAV-null *n* = 9; AAV-hTyr *n* = 11; Mann-Whitney test with Holm-Sidak’s correction; days: 1 (P = 0.999), 2 (P = 0.999), 3 (P = 0.945), 4 (P = 0.996), 5 (P = 0.999), 6 (P = 0.995), 7 (P = 0.995) and 8 (P = 0.821). **C-E)** Data are expressed as mean ± SEM. #P < 0.05 (between factors); **P < 0.01 (within factors AAV-null).

**Extended Data Fig. 6.**
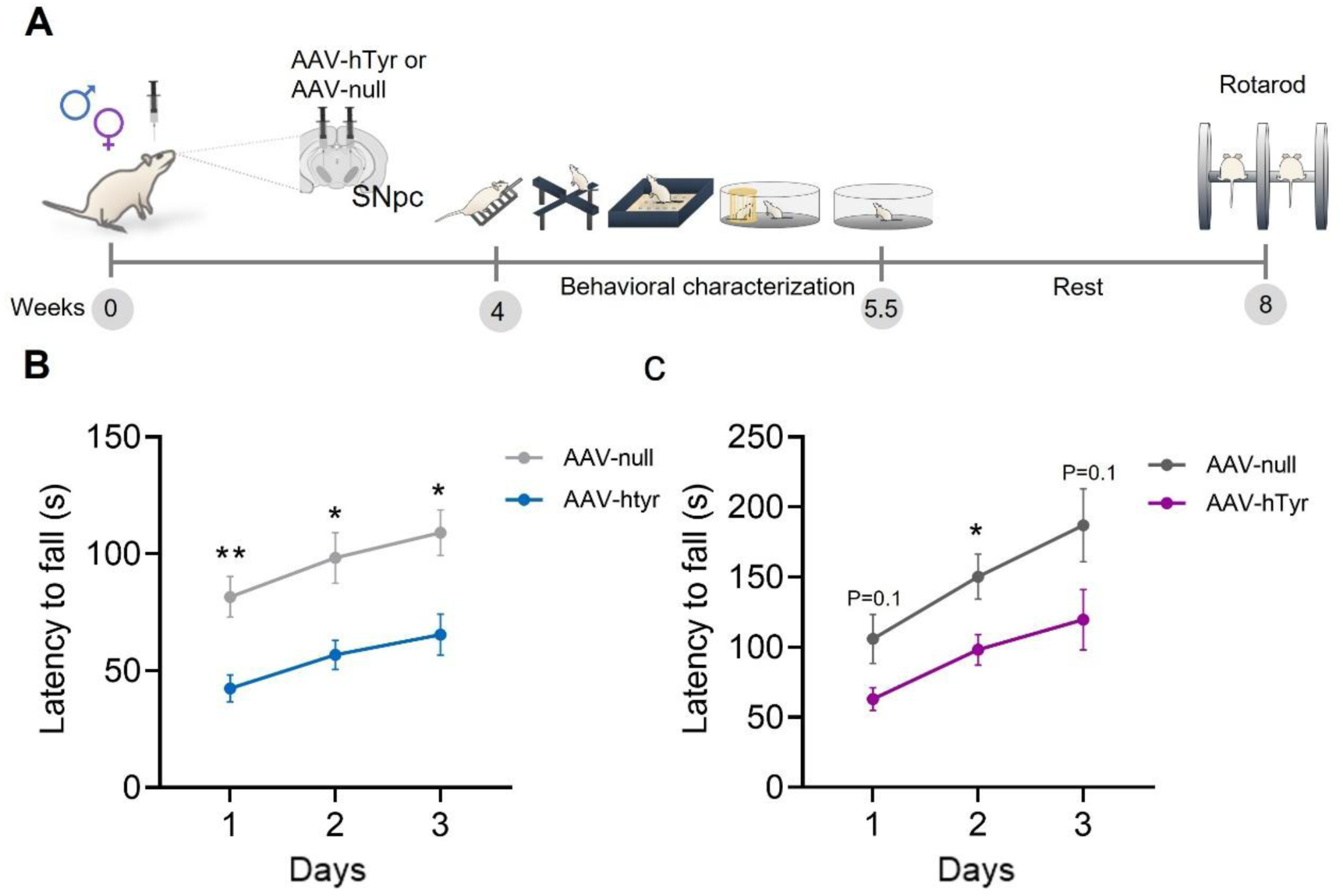
Sustained NM buildup impairs motor coordination irrespective of sex **A,** Experimental timeline and design used to evaluate rats’ motor coordination with the rotarod test at advanced stages of NM buildup (8 weeks post-AAV-hTyr injection). B, Plot of latency to fall from rotarod during three daily sessions by male AAV-null (*n* = 10) and AAV-hTyr-injected rats (*n* = 10), showing that AAV-hTyr-injected rats fall earlier than controls. RM two-way ANOVA followed by Sidak’s test, days 1 (P = 0.005), 2 (P = 0.015) and 3 (P = 0.011). **C,** Plot of latency to fall from rotarod across different sessions by female rats indicating that AAV-hTyr-injected rats (n = 9) fall earlier from rotarod compared to AAV-null (n = 10). Multiple Mann-Whitney test with Holm-Sidak correction, days 1 (P = 0.109), 2 (P = 0.030) and 3 (P = 0.109). **B-C,** Data are expressed as mean ± SEM. *P < 0.05.

